# ESCRT-I Inhibition Protects against NMDAR Hypofunction and Restores Synaptic Homeostasis in a Cellular Model of Schizophrenia

**DOI:** 10.64898/2026.09.07.749795

**Authors:** Mohamed Fouad Shalaby, Vincenzo Prato, Jacobo Elies Gomez, Nikita Gamper, Samantha Louise McLean, Sriharsha Kantamneni

## Abstract

N-methyl-D-aspartate receptor (NMDAR) hypofunction is a central pathophysiological mechanism in schizophrenia, yet direct NMDAR potentiation has shown limited clinical benefit, potentially because it fails to address deficits in receptor trafficking and surface stability. The endosomal sorting complexes required for transport (ESCRT) machinery governs the lysosomal fate of internalised membrane proteins, but its role in synaptic receptor homeostasis during glutamatergic dysfunction remains unclear. Phencyclidine (PCP), a non-competitive NMDAR antagonist, is widely used to model NMDAR hypofunction and schizophrenia-like phenotypes. Here, we show that sub-chronic PCP exposure induced persistent NMDAR hypofunction, impaired GABAergic transmission, and collapsed of excitation/inhibition (E/I) balance in primary hippocampal neurons. Genetic inhibition of the ESCRT-I component TSG101 prevented these deficits, enabled functional recovery following PCP washout, and increased surface expression of functional NMDARs and GABAA receptors without altering receptor pharmacology or biophysical properties. At the network level, TSG101 knockdown restored inhibitory tone, normalised excitatory activity, and stabilised E/I balance. Moreover, TSG101 knockdown rescued PCP-induced reductions in PSD-95 and BDNF, restored ERK1/2 signalling, and re-established activity-dependent nuclear translocation of Fos-like (Fos-L), consistent with reactivation of transcriptional plasticity. These findings identify ESCRT-I as a regulator of synaptic stability, positioning endosomal sorting pathways as a potential therapeutic target for schizophrenia.

## 1 – Introduction

The glutamate hypothesis of schizophrenia posits that reduced N-methyl-D-aspartate receptor (NMDAR) function contributes to the core cognitive and negative symptoms of the disorder [1, 2]. This hypothesis is supported by converging lines of evidence. These include: NMDAR antagonists such as phencyclidine (PCP) and ketamine replicate schizophrenia-like symptoms in healthy individuals and exacerbate symptoms in patients [3, 4]; reduced NMDAR expression and altered subunit composition in the post-mortem brains of people with schizophrenia [5, 6]; and convergence of genetic risk factors on synaptic proteins that regulate NMDAR trafficking and function [7, 8]. Collectively, these observations strongly support the role of NMDAR hypofunction as a core pathophysiological mechanism.

Cortical disinhibition is a key mechanistic consequence of NMDAR hypofunction. NMDARs located on parvalbumin-positive inhibitory GABAergic interneurons are particularly sensitive to blockade, leading to reduced interneuron firing, decreased GABA release, and subsequent hyperexcitability of pyramidal neurons [9, 10]. Consequent disruption of excitation/inhibition (E/I) balance is thought to underlie aberrant network oscillations, impaired sensory gating, and the cognitive dysfunction observed in schizophrenia [11, 12]. Strategies that increase glutamatergic tone without concomitantly strengthening inhibitory circuits risk exacerbating network instability [13]. Therefore, restoring E/I balance, rather than simply augmenting excitatory drive, could represent an effective therapeutic intervention.

Consistent with this hypothesis, studies aimed at directly potentiating NMDARs have yielded disappointing results in clinical trials. Glycine-site agonists, D-serine, and NMDAR positive allosteric modulators have shown only modest or inconsistent efficacy [14, 15]. These failures may reflect the inability of direct receptor potentiation strategies to correct underlying deficits in receptor trafficking and surface stability, both of which are critical for maintaining functional receptor pools at synapses [16, 17]. This raises the possibility that targeting the intracellular machinery governing receptor surface availability may provide a more effective therapeutic approach.

Surface expression of synaptic receptors is a tightly regulated process governed by cycles of exocytosis, lateral diffusion, endocytosis, and either lysosomal degradation or recycling back to the plasma membrane [16, 18]. NMDARs undergo activity-dependent internalisation via clathrin-mediated endocytosis, and the balance between receptor recycling and degradation determines the size of the functional receptor pool at the cell surface. [19, 20].

Interestingly the NMDAR hypofunction hypothesis of schizophrenia proposes that NMDAR hypofunction is preferentially expressed on fast-spiking GABAergic interneurons, where it disrupts interneuron excitability and disinhibits downstream pyramidal neurons, producing a net increase in cortical excitatory drive and collapse of E/I balance [21]. In schizophrenia, converging evidence points to dysregulation of the trafficking equilibrium underlying this circuit-level deficit. Reduced expression of postsynaptic density scaffolding proteins, such as PSD-95, impairs NMDAR anchoring at both excitatory and inhibitory synapses, with particular functional consequences at interneuron synapses where NMDAR loss disproportionately impairs interneuron recruitment [22, 23]. Enhanced ubiquitin-proteasome activity accelerates receptor turnover [24] and genetic variants affecting endosomal sorting pathways have been identified in genome-wide association studies [7, 8]. These observations are consistent with aberrant receptor degradation, rather than reduced receptor synthesis, as a primary driver of NMDAR hypofunction. Thus, interventions aimed at promoting surface retention, particularly on interneurons, could have therapeutic value.

The endosomal sorting complexes required for transport (ESCRT) machinery governs the lysosomal trafficking and degradation of internalised membrane proteins [25, 26]. Through the sequential actions of the ESCRT-0, -I, -II, and -III complexes, ubiquitinated cargo is sorted into the intraluminal vesicles of multivesicular bodies, thereby regulating receptor turnover and surface availability [27, 28]. The ESCRT-I component TSG101 (Tumour Susceptibility Gene 101) recognises ubiquitinated cargo and initiates this degradative sorting, whereas the AAA-ATPase VPS4a catalyses ESCRT-III disassembly to enable membrane scission and component recycling [29, 30].

ESCRT dysfunction has been implicated in neurodegenerative disorders including frontotemporal dementia [31, 32], and transcriptomic analyses of schizophrenia postmortem tissue reveal altered expression of multiple ESCRT complex genes, including TSG101, Hepatocyte Growth Factor-Regulated Tyrosine Kinase Substrate (HGS), and Charged Multivesicular Body Protein 2B (CHMP2B), suggesting that impaired endosomal sorting may also contribute to the synaptic pathology of neuropsychiatric disorders [7, 8]. However, the functional consequences of ESCRT perturbation on glutamatergic and GABAergic receptor homeostasis, and its potential as a therapeutic target, remain unexplored.

For this study, we employed a sub-chronic PCP treatment regimen followed by a washout period, rather than an acute challenge. In primary neuronal cultures, this approach allows us to distinguish between the immediate, direct effects of receptor blockade and the more enduring, persistent synaptic adaptations that arise from sustained NMDAR hypofunction [33]. Acute PCP exposure primarily reflects the direct pharmacological inhibition of the receptor. In contrast, sub-chronic exposure, by allowing a period of drug-free recovery, permits the study of persistent changes in receptor trafficking, network excitability, and synaptic protein expression that are thought to model the core, enduring pathophysiology of schizophrenia more closely [33, 34].

Here, we used sub-chronic phencyclidine (PCP) treatment in primary cortical neurons as a cellular model of schizophrenia-associated glutamatergic hypofunction to investigate if targeting ESCRT-I-dependent receptor trafficking protects against persistent synaptic deficits. Specifically, we examined whether genetic inhibition of TSG101 alters the surface availability of NMDARs and GABA_A_ receptors, modifies the functional consequences of PCP-induced E/I imbalance, and engages downstream molecular pathways of synaptic plasticity. By combining whole-cell patch-clamp electrophysiology, calcium imaging, surface biotinylation, and immunocytochemistry, we show that TSG101 inhibition modifies receptor surface expression and preserves synaptic and network function following PCP-induced NMDAR hypofunction, accom-panied by recovery of molecular markers associated with synaptic plasticity.

## 2 – Materials and Methods

### 2.1 – Primary neuronal culture

All animal procedures were approved by the University of Bradford Animal Welfare and Ethical Review Body and complied with UK Home Office regulations under the Animal (Scientific Procedures) Act 1986. Primary cortical and hippocampal neurons were prepared from embryonic day 18 (E18) Sprague-Dawley rat embryos as previously described [25]. Briefly, embryos were rapidly dissected in ice-cold Hank’s Balanced Salt Solution (HBSS; Gibco) calcium- and magnesium-free. Cortices and hippocampi were isolated, pooled, and enzymatically digested with 0.25% trypsin-EDTA (Gibco) for 15 min at 37°C. Digestion was terminated by adding heat-inactivated horse serum (HS; Gibco). Tissue was triturated in HBSS containing 10 µg/mL DNase I (Sigma-Aldrich) and 1 mg/mL soybean trypsin inhibitor (Sigma-Aldrich) using fire-polished Pasteur pipettes of decreasing bore size. Cells were passed through a 70 µm cell strainer and plated onto poly-D-lysine (0.1 mg/mL; Sigma-Aldrich) and laminin (2 µg/mL; Sigma-Aldrich)-coated coverslips or culture plates at a density of 75,000 cells/cm². Neurons were maintained in Neurobasal Plus medium (Gibco) supplemented with B-27 Plus (Gibco), 2 mM GlutaMAX (Gibco), 5% HS, 10 U/mL penicillin, and 10 µg/mL streptomycin (Gibco) at 37°C in a 5% CO₂ humidified incubator. Half of the culture medium was replaced every 3–4 days. All experiments were performed on cultures at days in vitro (DIV) 12–14, a stage of mature synaptic connectivity.

### 2.2 – Lentiviral Production and Neuronal Transduction

Lentiviral constructs were obtained from VectorBuilder and included: scrambled shRNA control (VB200120-1037smy), shRNA targeting rat TSG101 (shTSG101; VB900077-7127vsc), dominant-negative VPS4a (DN-VPS4a; E228Q mutant; VB240813-1403ubj), wild-type rat TSG101 overexpression (rTSG101; VB240813-1424gea), wild-type rat VPS4a overexpression (rVPS4a; VB240809-1242ajb), and EGFP/mCherry reporter controls (VB240813-1423gnb). All constructs were sequence-verified prior to use.

Lentiviral particles were generated in HEK293T cells (ATCC CRL-3216) using a three-plasmid second-generation packaging system. Cells were seeded in 10 cm dishes and co-transfected at 70–80% confluence using Lipofectamine 3000 (Thermo Fisher) with the transfer vector (gene of interest), PLP1 (gag/pol), PLP2 (rev), and pVSV-G (envelope glycoprotein) at a mass ratio of 45:20:15:20, respectively. DNA– lipid complexes were prepared in Opti-MEM (Gibco) and applied to cells for 6 h before medium replacement. Viral supernatants were collected at 48 h and 72 h post-transfection, clarified by centrifugation (500 × g, 5 min), filtrated through 0.45 µm membranes, aliquoted, and stored at −80°C. Functional viral titres were estimated by serial dilution and quantification of transduction efficiency in HEK293T cells.

At DIV7, primary neurons were transduced with lentiviral particles at equivalent titres (multiplicity of infection, 5–10) in the presence of 8 µg/mL polybrene (Sigma-Aldrich). After 12–16 h, the medium was replaced with fresh Neurobasal Plus medium. All experiments were performed at DIV12–14 (5–7 days post-transduction), allowing sufficient time for stable transgene expression and efficient gene knockdown.

### 2.3 – Surface Biotinylation Assays

Cell-surface receptor expression was quantified using EZ-Link Sulfo-NHS-SS-Biotin (Thermo Fisher) following established methods for membrane protein isolation [35]. Cells were washed with ice-cold PBS and incubated with 0.5 mg/mL sulfo-NHS-SS-biotin in HBSS (Gibco) for 30 min at 4°C with gentle agitation. Excess reagent was quenched with 100 mM glycine in PBS for 10 min at 4°C. Cell lysates were prepared in RIPA buffer supplemented with protease inhibitor cocktail (Roche), and biotinylated proteins were isolated using Pierce™ High-Capacity Streptavidin Agarose (Thermo Fisher). Surface and total protein fractions were subjected to SDS-PAGE and immunoblotting. GAPDH and/or β-actin were used as loading controls for total lysates and as negative controls to confirm the absence of cytosolic contamination in surface fractions.

#### Immunoblotting

Proteins were resolved on 8% SDS PAGE gels (Bio Rad, Hercules, CA, USA) and transferred to PVDF membranes (Millipore, USA). Membranes were blocked in 5% BSA (Sigma-Aldrich) in Tris-buffered saline with 0.1% Tween-20 (TBST) for 1 h at room temperature and incubated overnight at 4°C with primary antibodies against GluN1 (mouse, clone 1A4B10, Synaptic Systems), GluN2A (rabbit, AB1555P, Synaptic Systems), GABA_A_RG2 (rabbit, AB_223344, Abcam), GABA_B_R1 (guinea pig, AB_11234, Abcam), TSG101 (rabbit, Ab30871, Abcam), VPS4a (mouse, SAB4200215, Sigma-Aldrich), EGFR (rabbit, AB52894, Abcam), β-actin (mouse, AC-15, Sigma-Aldrich), and GAPDH (Rabbit, EPR16891, Abcam). Fluorescent secondary antibodies (IRDye 680RD anti-mouse IgG, 925-68070; IRDye 800CW anti-rabbit IgG, 925-32211; LI-COR Biosciences) were applied for 1 h at room temperature. Membranes were imaged on a LI-COR Odyssey DLx system and band intensities were quantified using ImageJ (Fiji, version 2.16.0/1.54p).

### 2.4 – Sub-Chronic Phencyclidine Treatment

Phencyclidine hydrochloride (PCP; Sigma-Aldrich, P3020) was supplied as a 1 mg/mL (approximately 2.7 mM) aqueous solution. For sub-chronic treatment, at DIV 21, neurons were incubated with 10 µM PCP in culture medium for 7 days, with medium refreshed every 2–3 days to maintain drug concentration. Control cultures received equivalent volumes of vehicle (PBS) under identical conditions. Following treatment, drug-containing medium was completely aspirated, and neurons were washed three times with pre-warmed Neurobasal Plus medium. Cells were then incubated in fresh, drug-free culture medium for 48 h prior to electrophysiological, biochemical, or imaging analyses to model persistent post-washout deficits rather than acute receptor blockade.

### 2.5 – Whole-Cell Patch-Clamp Electrophysiology

Whole-cell voltage-clamp recordings were performed at room temperature (22–24°C) on visually identified pyramidal-like neurons using an multiclamp 700B amplifier and Digidata 1550B digitiser (Molecular Devices). Data were acquired and analysed using pClamp 11 software (Molecular Devices). Recording electrodes (3–5 MΩ) were pulled from borosilicate glass capillaries (World Precision Instruments) using a horizontal micropipette puller (Sutter Instruments). A fine-bore perfusion tube positioned 0.2 cm above the recorded cell at a controlled rapid flow rate was used to ensure localised, minimally diluted drug delivery. Series resistance was monitored throughout each recording; cells were excluded if series resistance exceeded 20 MΩ or changed by more than 20% during recording.

For NMDAR current recordings, the external solution contained (in mM): 140 NaCl, 5 KCl, 2 CaCl₂, 10 HEPES, and 10 glucose, with MgCl₂ omitted to relieve voltage-dependent Mg²⁺ block (pH 7.4, 310–320 mOsm). The internal solution contained (in mM): 135 CsMeSO₄, 8 NaCl, 10 HEPES, 0.3 EGTA, 4 Mg-ATP, 0.3 Na-GTP, and 20 tetraethylammonium (TEA; pH 7.3, 290–300 mOsm). AMPA and kainate receptor-mediated currents were blocked by the inclusion of 10 µM CNQX (Tocris), and action potentials suppressed with 1 µM tetrodotoxin (TTX; Tocris). NMDAR currents were evoked by a 2 s bath application of 30 µM NMDA (Tocris) and 10 µM glycine (Sigma-Aldrich) at holding potentials ranging from −60 to +60 mV. Peak current amplitudes were measured at each holding potential and normalised to the current recorded at +60 mV for current–voltage (I-V) analysis [36–38].

For AMPAR current recordings, neurons were voltage-clamped at −60 mV in external solution containing 1 mM MgCl₂. AMPA receptor-mediated currents were evoked by a 2 s bath application of 100 µM AMPA (Tocris) [39]. For NMDA/AMPA ratio analysis, NMDAR currents were evoked by a 2 s co-application of 30 µM NMDA and 10 µM glycine in Mg²⁺-free external solution. NMDAR current amplitudes were measured 50 ms after agonist application to ensure near-complete decay of the fast AMPA receptor-mediated component.

For GABA_A_ receptor recordings, the external solution contained (in mM): 140 NaCl, 5 KCl, 2 CaCl₂, 1 MgCl₂, 10 HEPES, and 10 glucose (pH 7.4). The internal solution contained (in mM): 135 CsCl, 8 NaCl, 10 HEPES, 0.3 EGTA, 4 Mg-ATP, and 0.3 Na-GTP (pH 7.3). GABA(A) receptor-mediated currents were evoked by a 2 s bath application of GABA (0.01–100 µM; Sigma-Aldrich) at a holding potential of −60 mV [40–42]. Concentration-response curves were fitted to the Hill equation, yielding EC50 values of 0.056 µM (Control/SCR) and 0.059 µM (shTSG101) (see results). These values are consistent with high-affinity extrasynaptic GABA_A_ receptors previously described in hippocampal neurons [52][42, 43]. For phenobarbital potentiation experiments, submaximal currents were evoked using the EC_50_ concentration of GABA (0.06 µM) in the absence or presence of phenobarbital (100 µM; Sigma-Aldrich). To assess bicuculline-mediated inhibition, currents were evoked by 1 µM GABA in the presence or absence of bicuculline (10 µM; Tocris).

For spontaneous excitatory postsynaptic currents (sEPSCs) and spontaneous inhibitory postsynaptic currents (sIPSC) recordings, primary hippocampal neurons were visually identified based on morphological criteria, both sEPSCs and sIPSCs were recorded from the same population of pyramidal-like neurons, identified by their large, triangular somata and prominent apical dendrites. This approach allowed assessment of both excitatory and inhibitory inputs converging onto the same cell type. All spontaneous recordings were performed at a holding potential of −60 mV in external solution containing (in mM): 140 NaCl, 5 KCl, 2 CaCl₂, 1 MgCl₂, 10 HEPES, and 10 glucose (pH 7.4) [44].

sEPSCs were recorded using the CsMeSO₄-based internal solution in an antagonist-free external solution. sIPSCs were recorded at using the CsCl-based internal solution and an external solution supplemented with 10 µM CNQX and 50 µM D-AP5 (Tocris) to block glutamatergic transmission [45, 46]. Continuous recordings were acquired for 5 min per cell. Synaptic events were detected using a threshold-based algorithm in Clampfit (Molecular Devices), with the detection threshold set at three times the baseline root mean square (RMS) noise and subsequently verified by visual inspection. Events frequency and amplitude were quantified for each cell. The inhibition/excitation (I/E) ratio was calculated for each neuron as the ratio of sIPSC frequency to sEPSC frequency.

### 2.6 – Calcium Imaging

Intracellular calcium dynamics were assessed using the fluorescent calcium indicators Fluo-4 AM ([Thermo Fisher, UK]) and Rhod-2 AM (Abcam). Neurons plated in black-walled, clear-bottom 96-well plates (Corning) were loaded with 5 µM Fluo-4 AM or 5 µM Rhod-2 AM in HBSS containing 20 mM HEPES (pH 7.4), 0.02% Pluronic F-127 (Thermo Fisher), and 0.5% probenecid (Sigma-Aldrich) for 45 min at 37°C in the dark. Cells were washed twice with dye-free buffer and allowed to de-esterify for 20 min at 37°C [47].

Fluorescence was recorded using a FlexStation 3 Multi-Mode Microplate Reader (Molecular Devices). For Fluo-4 AM experiments, excitation and emission wavelengths were set at 485 and 520 nm, respectively. For Rhod-2 AM experiments, excitation and emission wavelengths were set at 552 and 581 nm, respectively. Baseline fluorescence was recorded for 30 s, followed by automated addition of agonist (NMDA, 3.125–100 µM, in the presence of 10 µM glycine; AMPA, 0.01–100 µM; Tocris) or vehicle. For antagonist experiments, PCP (0.1–30 µM; Sigma-Aldrich), CNQX (0.3–30 µM; Tocris), or NBQX (0.3–30 µM; Tocris) was pre-incubated for 160 s prior to agonist addition. Fluorescence was recorded for a total duration of 180– 300 s. Data were analysed using SoftMax Pro software (Molecular Devices). Calcium transients were quantified as ΔF/F₀ = (Fmax − F₀)/F₀, where F₀ represents baseline fluorescence and Fmax represents peak fluorescence. Area under the curve (AUC) was calculated using the trapezoidal method: AUC = (Fmax∼ + Fsteady)/2 × (T_end – t_peak) + (F₀ + Fmax)/2 × (t_peak − t₀). Dose-response curves were fitted by nonlinear regression analysis in GraphPad Prism.

### 2.7 – Immunocytochemistry

Cell-surface receptor localisation was assessed using non permeabilised immunocytochemistry, as previously described for extracellular epitope detection (49). For surface receptor staining, neurons were fixed with 4% paraformaldehyde (PFA) in PBS for 15 min at room temperature without permeabilisation. Cells were blocked in 5% normal goat serum (NGS) in PBS for 30 min and incubated overnight at 4°C with primary antibodies recognising extracellular epitopes of GluN1 (1:200; Synaptic Systems, 1A4B10), GluN2A (1:200; Synaptic Systems, AB1555P), GABRG2 (1:200; Abcam, AB223344), or GABABR1 (1:200; Abcam, AB11234). Following washing, cells were incubated for 1 h at room temperature with Alexa Fluor 488- or 594-conjugated secondary antibodies (1:500; Jackson ImmunoResearch). Nuclei were counterstained with DAPI (1 µg/mL; Thermo Fisher).

For intracellular staining, neurons were permeabilised with 0.1% Triton X-100 in PBS for 10 min following fixation, blocked in 5% NGS, and incubated overnight at 4°C with primary antibodies against PSD-95 (1:500; Synaptic Systems), BDNF (1:500; Abcam), pERK1/2 (1:200; Cell Signalling Technology), or Fos-L (1:200; Sigma-Aldrich). Secondary antibody incubation and nuclear counterstaining were performed as described above.

Images were acquired using a Zeiss Axiovert 200 M confocal microscope equipped with a 63× oil-immersion objective (NA 1.3) or a Nikon Ti-S inverted fluorescence microscope. Laser power, detector gain, and offset settings were kept constant across all experimental conditions. Fluorescence intensity, puncta density, and colocalisation were quantified using ImageJ. For neuronal analyses, 6–8 neurons per condition were analysed from at least three independent culture preparations.

### 2.8 – Statistical Analysis

All data are presented as mean ± standard error of the mean (SEM). Statistical analyses were performed using GraphPad Prism 9 (GraphPad Software). Data normality was assessed using the Shapiro-Wilk test. Comparisons between two groups were performed using unpaired, two-tailed Student’s t-test for normally distributed data or Mann-Whitney U test for non-normally distributed data. Comparisons across multiple groups were conducted using one-way or two-way analysis of variance (ANOVA), followed by Tukey’s or Dunnett’s post-hoc tests for multiple comparisons, as appropriate.

For electrophysiological data, the number of cells (n) represents individual neurons recorded from at least 3 independent culture preparations. For biochemical experiments, n represents independent experiments performed on separate culture preparations. P values of less than 0.05 were considered statistically significant and denoted as follows: p < 0.05 (*), p < 0.01 (**), p < 0.001 (***), and p < 0.0001 (****).

## 3 – Results

### 3.1 – Interfering with ESCRT Function Differentially Regulates Neurotransmitter Receptor Surface Localisation in Primary Neurons

To determine whether ESCRT-dependent trafficking regulates the surface availability of endogenous neurotransmitter receptors we examined receptor localisation in rat primary hippocampal cultures following genetic perturbation of ESCRT-I and ESCRT-III components.

TSG101 knockdown significantly increased the surface-to-total ratios of the NMDA receptor subunits GluN1 and GluN2A in primary neurons, as determined by surface biotinylation assays (Fig. 1A–C). Immunocytochemical analyses performed under non-permeabilised conditions confirmed enhanced dendritic surface localisation of both subunits following shTSG101 expression (Fig. 1N–O). To determine whether these effects were specific to ESCRT-I inhibition, we disrupted ESCRT-III function using a dominant-negative mutant of the AAA-ATPase VPS4a (DN-VPS4a; E228Q), which traps ESCRT-III complexes in a non-functional state. DN-VPS4a expression increased the surface abundance of GluN1 and GluN2A (Fig. 1D-F), suggesting that ESCRT-III disruption also promotes NMDAR surface accumulation.

**Figure 1:**
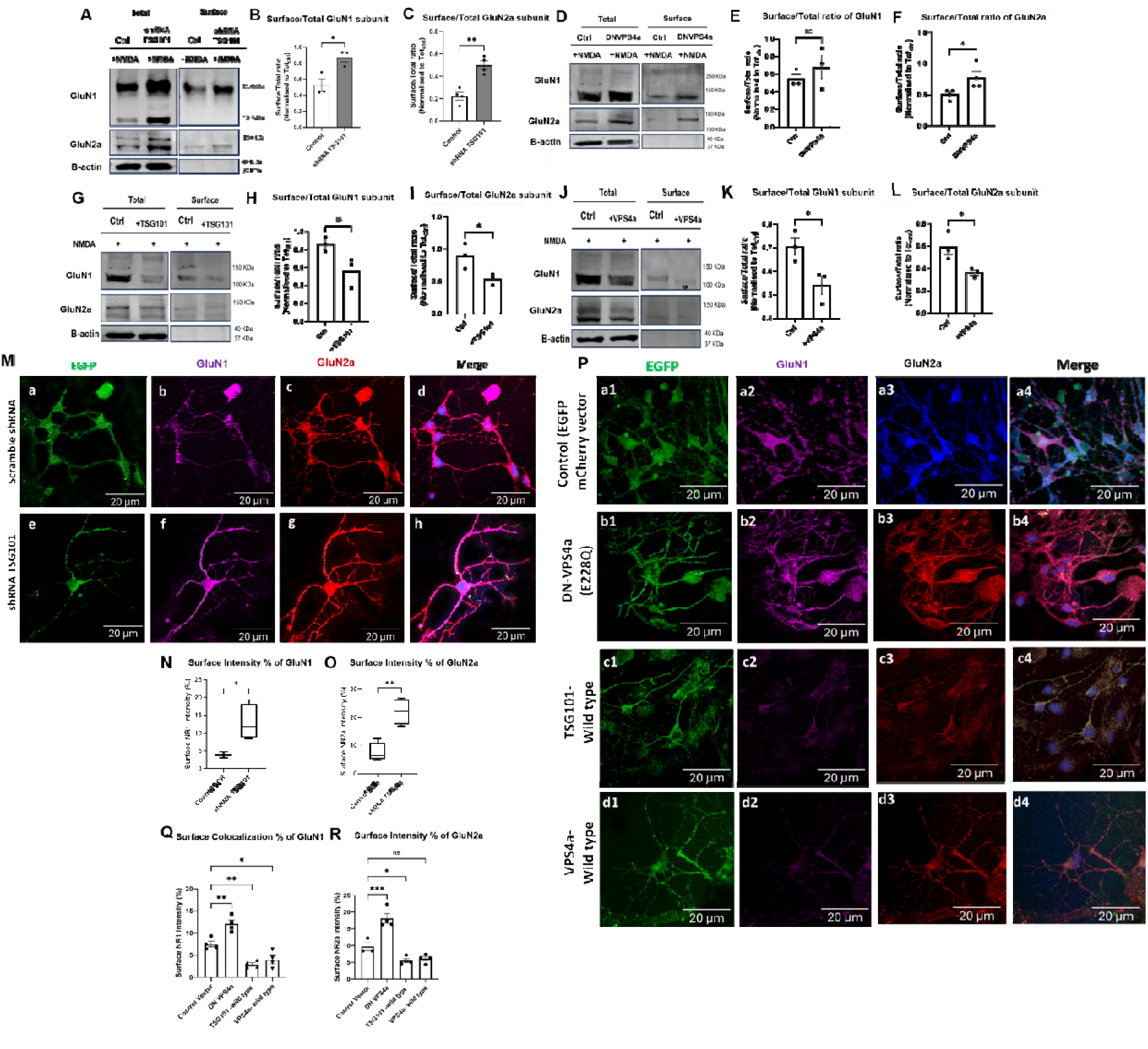
ESCRT pathway perturbation regulates surface NMDAR expression in primary neurons. (A–C) Effect of shTSG101-mediated knockdown on NMDAR subunit surface expression. (A) Representative surface biotinylation immunoblots for GluN1 and GluN2a (total and surface fractions); β-actin served as a loading control. (B–C) Quantification of the surface-to-total ratios, showing a significant increase in surface GluN1 and GluN2a following TSG101 knockdown compared to control neurons. (D–F) Effect of dominant-negative VPS4a (DN-VPS4a) on NMDAR trafficking. (D) Representative immunoblots of total and surface fractions. (E–F) Quantification of surface-to-total ratios, showing a significant reduction in surface expression in DN-VPS4a-expressing neurons. (G–I) Effect of TSG101 overexpression (+TSG101). (G) Representative immunoblots. (H–I) Quantification of surface-to-total ratios showing reduced surface receptor abundance, suggesting enhanced internalization and/or degradation. (J–L) Effect of wild-type VPS4a overexpression (+VPS4a). (J) Representative immunoblots. (K–L) Quantification of surface-to-total ratios, demonstrating decreased receptor surface abundance. (M–O) Immunocytochemical analysis of surface NMDARs under non-permeabilised staining conditions in primary cortical neurons. (M) Representative confocal images of control (a–d) and shTSG101-transduced (e–h) neurons (EGFP, green; surface GluN1, magenta; surface GluN2a, red). Merged images show increased receptor localisation at neuronal processes following TSG101 knockdown. (N–O) Quantification of surface fluorescence intensity. (P–R) Representative confocal images and quantification of NMDAR surface localization following ESCRT modulation. (P) Images of control (a1–a4), DN-VPS4a-expressing (b1–b4), TSG101-overexpressing (c1–c4), and VPS4a-overexpressing (d1–d4) neurons (EGFP, green; surface GluN1, magenta; surface GluN2a, red). Merged images demonstrate reduced surface localisation following ESCRT enhancement or manipulation of VPS4 ATPase activity. (Q–R) Quantification of surface GluN1 and GluN2a fluorescence intensity. Data are mean ± SEM (n = 3–5 independent experiments). Statistical significance was determined by one-way ANOVA followed by Tukey’s post-hoc test. *P < 0.05, **P < 0.01, ***P < 0.001, ****P < 0.0001; ns, not significant. Scale bars = 20 μm.

Comparative immunofluorescence demonstrated that receptor distribution patterns differed qualitatively between shTSG101 and DN-VPS4a. A diffuse, punctate labelling pattern across dendritic shafts and spines was observed with shTSG101, consistent with enhanced receptor recycling and surface delivery. DN-VPS4a resulted in more punctate, perinuclear aggregates, potentially reflecting receptor trapping in late endosomal compartments (Fig. 1Q–R). These findings suggest distinct mechanistic consequences of ESCRT-I versus ESCRT-III disruption on glutamate receptor trafficking.

Subsequently, we investigated the effects of wild-type TSG101 and VPS4a. Both TSG101 (Fig. 1G–I) and VPS4a (Fig. 1J–L) overexpression significantly reduced surface NMDAR levels, consistent with enhanced ESCRT-I-mediated receptor degradation. Collectively, these data demonstrate that ESCRT activity regulates the steady-state surface pool of NMDARs, with ESCRT-I and ESCRT-III exerting overlapping but distinct regulatory roles.

TSG101 knockdown also significantly increased the surface expression of GABA receptor subunits. Surface biotinylation assays revealed significantly increased surface-to-total ratios of the GABA_A_ receptor gamma-2 subunit (GABA_A_RG2) and the GABA_B_ receptor subunit 1 (GABA_B_R1) in shTSG101-expressing neurons (Fig. 2A– C). Immunofluorescence analysis confirmed correspondingly higher dendritic surface expression of both receptor subunits (Fig. 2Q–R). In neurons expressing DN-VPS4a, surface GABA_A_RG2 levels were similarly elevated (Fig. 2D–E), whereas GABA_B_R1 surface expression remained unchanged relative to scrambled controls (Fig. 2F). These findings suggest that GABA_B_R trafficking is preferentially sensitive to ESCRT-I rather than ESCRT-III perturbation.

**Figure 2.**
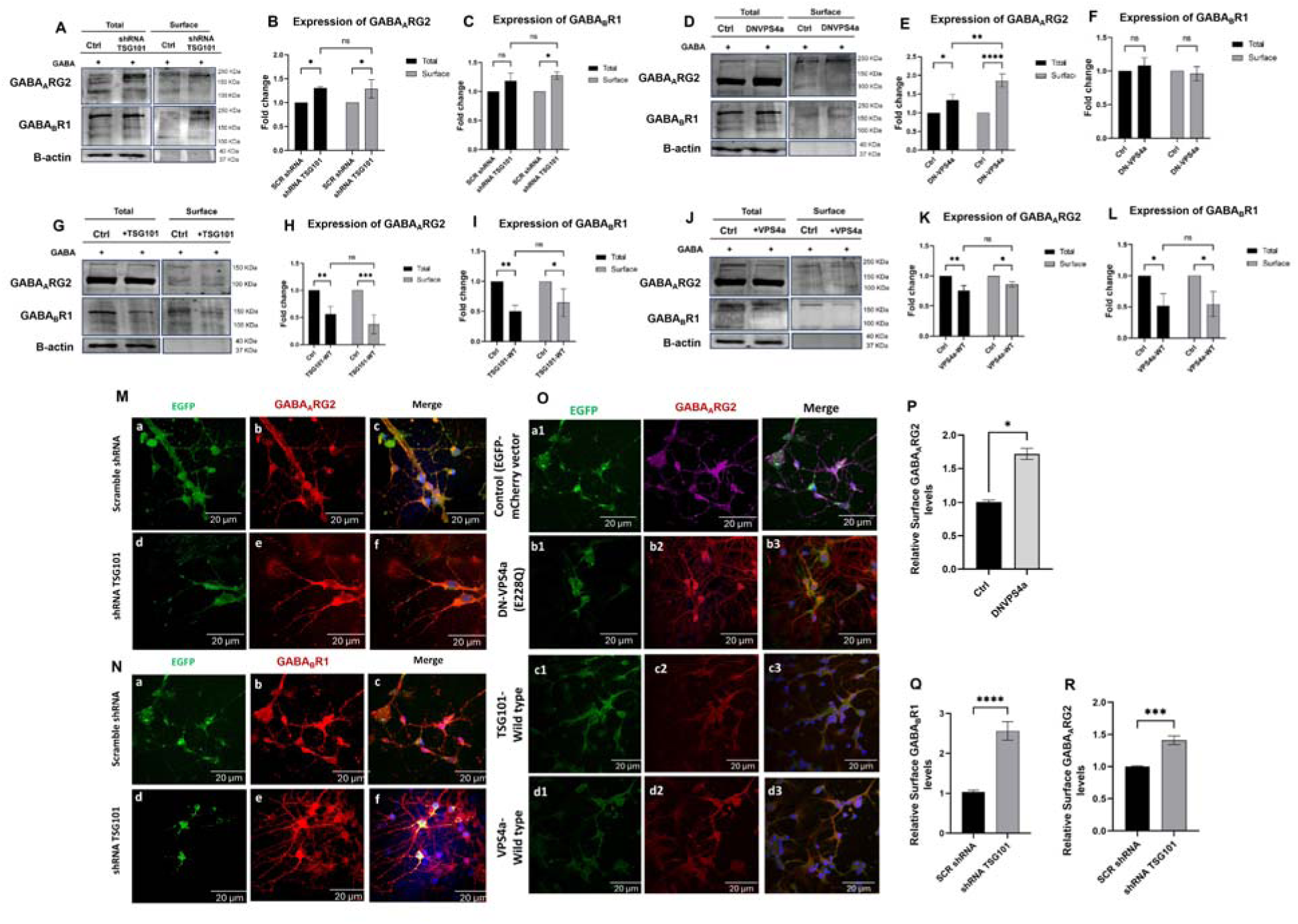
ESCRT-dependent regulation of GABA_A_ and GABA_B_ receptor subunits in primary cortical neurons. (A–C) Effect of TSG101 knockdown (shTSG101) on GABA_A_ and GABA_B_ receptor trafficking. (A) Representative surface biotinylation immunoblots for GABA_A_ receptor γ2 (GABA_A_RG2) and GABA_B_ R1/2 subunits (total and surface fractions); β-actin served as a loading control. (B–C) Quantification showing a significant increase in surface levels of both γ2 and R1/2 subunits following TSG101 knockdown, with total receptor abundance remaining largely unchanged. (D–F) Effect of dominant-negative VPS4a (DN-VPS4a) expression. (D) Representative immunoblots. (E–F) Quantification showing enhanced surface accumulation of the γ2 subunit, whereas R1/2 surface expression remained comparable to controls. (G–I) Effect of TSG101 overexpression (+TSG101). (G) Representative immunoblots. (H–I) Quantification demonstrating reduced surface expression of both subunits, consistent with enhanced ESCRT-mediated receptor sorting. (J–L) Effect of VPS4a overexpression (+VPS4a) (J) Representative immunoblots. (K–L) Quantitative analysis showing decreased surface receptor levels relative to controls. (M–N) Immunocytochemical analysis of surface receptors following TSG101 knockdown. Representative confocal images of control and shTSG101-transduced neurons (EGFP, green) stained for surface GABA_A_RG2 (M) or surface GABA_B_ R1 (N). Merged images demonstrate increased receptor localisation at the neuronal surface following TSG101 depletion. (O) Confocal analysis of GABA_A_RG2 surface localization following ESCRT modulation. Images show control, DN-VPS4a, TSG101-overexpressing, and VPS4a-overexpressing neurons (EGFP, green; GABA_A_RG2, magenta). Merged images illustrate increased surface accumulation following ESCRT inhibition and reduced surface localisation following ESCRT enhancement. (P–R) Quantification of surface immunofluorescence intensity for GABA_A_RG2 (P) and GABABR1 (Q) in control versus shTSG101 neurons, and (R) GABA_A_RG2 surface expression following broader ESCRT modulation. Data demonstrate increased receptor surface localisation following TSG101 depletion and reduced localisation following ESCRT enhancement. Data are mean ± SEM (n = 3–5 independent experiments). Statistical significance was determined by one-way ANOVA with Tukey’s post-hoc test or unpaired Student’s t-test. *P < 0.05, **P < 0.01, ***P < 0.001, ****P < 0.0001; ns, not significant. Scale bars = 20 μm.

Collectively, these results demonstrate that TSG101 knockdown mediated inhibition of ESCRT-I broadly enhances the surface availability of both excitatory and inhibitory receptor populations. ESCRT-III disruption through DN-VPS4a exerts more selective effects largely restricted to ionotropic receptor subunits.

### 3.2 – Sub-chronic PCP Treatment Induces Glutamatergic Hypofunction and Network Disinhibition

As a cellular model of NMDAR hypofunction, primary cortical and hippocampal neurons were exposed to sub-chronic PCP (10 µM, 7 days), followed by a 48-h washout period to enable the assessment of persistent post-treatment deficits rather than the acute effects of receptor blockade.

There was a marked impairment of NMDAR function following PCP treatment. Evoked NMDAR-mediated currents, recorded at +40 mV in Mg²⁺-free external solution containing CNQX to block AMPA receptors, were reduced by approximately 50% in PCP-treated neurons compared with vehicle controls (125.8 ± 13.5 pA vs. 250.5 ± 22.1 pA; n = 10 vehicle, n = 8 PCP, from 3 independent preparations; p < 0.001; Fig. 3A-B). Representative traces shown in Figure 3C, show the reduction was selective for NMDARs, as AMPAR-mediated currents were unchanged (Vehicle: -185.5 ± 16.2 pA; PCP: -172.3 ± 15.8 pA; n = 8 vehicle, n = 9 PCP, from 3 independent preparations; p > 0.05; Fig. 3D). Consequently, the NMDA/AMPA ratio, a measure of the relative contribution of NMDARs to synaptic transmission, was significantly reduced in PCP-treated neurons (0.28 ± 0.03 vs. 0.51 ± 0.04 in controls; n = 8 vehicle, n = 6 PCP, from 3 independent preparations; p < 0.01; Fig. 3E).

**Figure 3.**
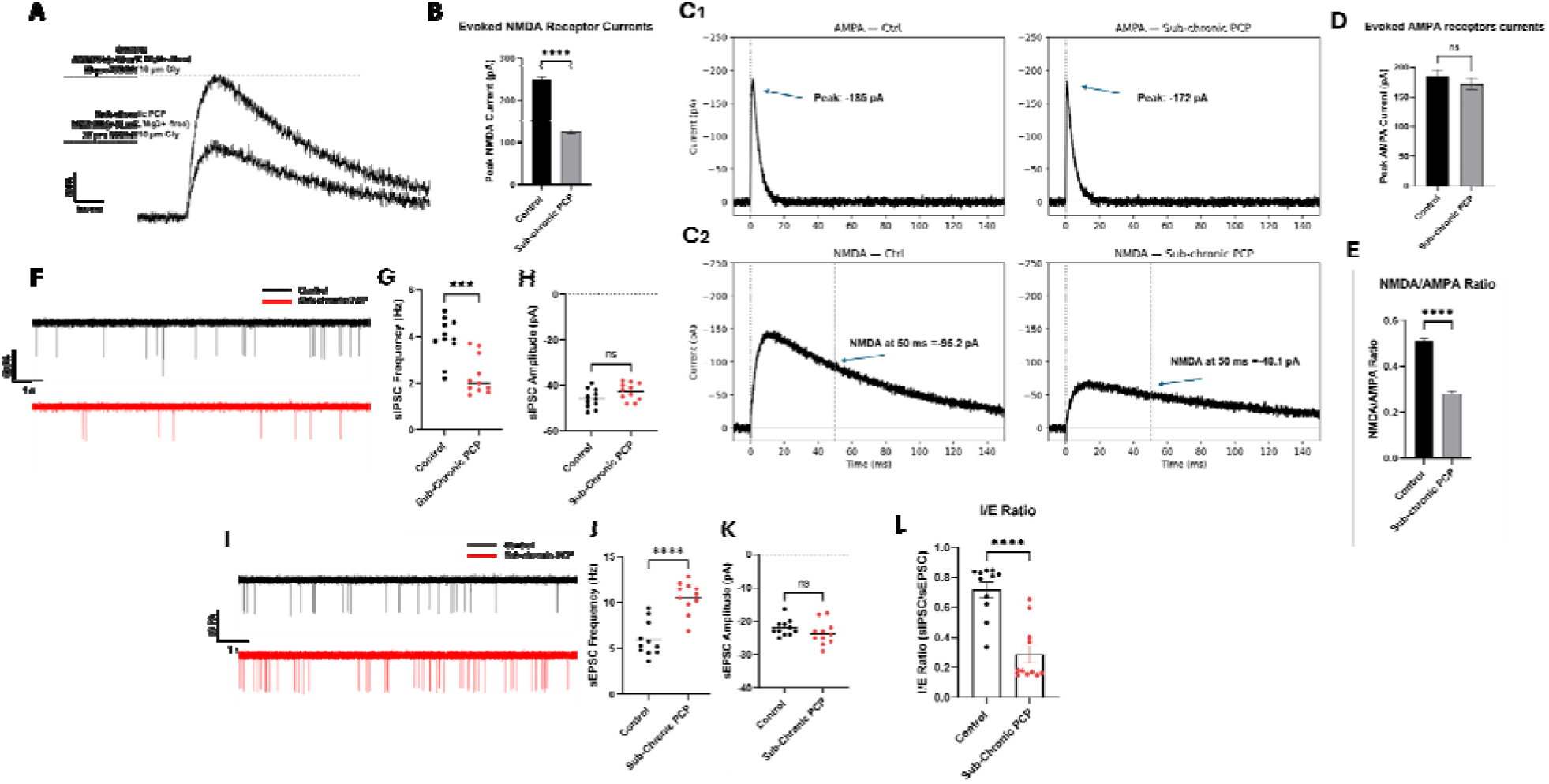
Sub-chronic PCP treatment induces glutamatergic hypofunction and network disinhibition. (A–B) NMDA receptor-mediated excitatory postsynaptic currents (EPSCs). (A) Representative whole-cell voltage-clamp recordings from control and sub-chronic PCP-treated neurons (recorded in Mg²⁺-free extracellular solution with 10 μM glycine). (B) Quantification of peak NMDA current amplitudes, showing a marked reduction in PCP-treated neurons. (C-E) NMDA/AMPA receptor current ratio. (C) Representative traces of AMPA and NMDA receptor-mediated EPSCs (AMPA currents measured at peak; NMDA currents measured 50 ms post-stimulation. (D) Quantification of peak AMPA current amplitudes, showing no significant difference. (E) Quantification of the NMDA/AMPA current ratio, showing a significant decrease in PCP-treated neurons, indicating impaired glutamatergic synaptic transmission. (F–H) Spontaneous inhibitory postsynaptic currents (sIPSCs). (F) Representative voltage-clamp traces. (G-H) Quantification of sIPSC frequency (G), which is significantly reduced, and amplitude (H), which remains unchanged, suggesting a predominantly presynaptic impairment in inhibitory transmission. (I–K) Spontaneous excitatory postsynaptic currents (sEPSCs). (I) Representative voltage-clamp traces. (J-K) Quantification of sEPSC frequency (J), showing a significant increase in PCP-treated neurons, and amplitude (K), which is unchanged. (L) Inhibitory/excitatory (I/E) ratio, calculated as the ratio of sIPSC to sEPSC frequency. PCP treatment significantly reduced the I/E ratio, indicating a shift toward network hyperexcitability and disrupted excitatory–inhibitory balance. Data are mean ± SEM (n = 8–15 cells from 3 independent preparations). Statistical significance was determined by unpaired Student’s t-tests. ****P < 0.0001; ***P < 0.001; ns, not significant.

To assess PCP treatment on inhibitory transmission, sIPSCs were recorded in the presence of CNQX and D-APV to isolate GABAergic events. PCP treatment significantly reduced sIPSC frequency from 4.2 ± 0.4 Hz to 1.9 ± 0.3 Hz (p < 0.001; Fig. 3F-G), whilst sIPSC amplitude remained unchanged (−45.8 ± 4.0 pA vs. −43.1 ± 3.9 pA; p > 0.05; Fig. 3H).

#### sEPSC amplitude reflects AMPAR function

Recordings of sEPSCs obtained in the absence of receptor blockers revealed a reciprocal increase in excitatory drive. PCP treatment more than doubled sEPSC frequency (11.3 ± 1.1 Hz vs. 5.1 ± 0.5 Hz; p < 0.001; Fig. 3I-J), with no significant change in sEPSC amplitude (−23.8 ± 2.2 pA vs. −22.5 ± 2.0 pA; p > 0.05; Fig. 3K).

Although sEPSCs were recorded in Mg²⁺-free solution to allow detection of both AMPA and NMDA receptor-mediated components, the fast rising/peak phase of each spontaneous event remains dominated by AMPA receptor-mediated currents, as NMDARs contribute primarily to the slower decaying component (Fig 3C). Thus, sEPSC peak amplitude predominantly reflects AMPAR-mediated current. This is consistent with our evoked recordings, where AMPAR-mediated currents were unchanged following PCP (Fig. 3C–E), whereas NMDAR-mediated currents, measured under conditions that isolate the NMDAR component (+40 mV, Mg²⁺-free), were significantly reduced (Fig. 3A–B).

We next calculated the I/E balance ratio (sIPSC frequency / sEPSC frequency) for each neuron sampled. PCP treatment significantly reduced the I/E ratio from 0.82 ± 0.08 in control neurons to 0.31 ± 0.04 (p < 0.001; Fig. 3L), confirming a shift towards a hyperexcitability indicative of disinhibited network state. Together, these findings indicate that the sub-chronic PCP washout paradigm represents a robust cellular model of persistent glutamatergic hypofunction and I/E imbalance relevant to schizophrenia pathophysiology.

### 3.3 – TSG101 Knockdown Enhances NMDA and GABA_A_ Receptor Function

To further investigate the mechanisms of ESCRT-I inhibition protection, we knocked down TSG101 in naïve neurons not treated with PCP to elucidate the mechanism by which ESCRT-I inhibition might confer resilience.

Whole-cell voltage-clamp recordings revealed that shTSG101 significantly increased evoked NMDAR current amplitudes across all tested holding potentials from -60 mV to +60 mV (Fig. 4A-B). The current-voltage (I-V) relationship revealed a significant increase in NMDAR cur-rent magnitude across all holding potentials following TSG101 knockdown (Fig. 4C), while normalised I-V curves overlapped (Fig. 4D), indicating that the voltage-dependence and rectification properties of NMDARs were unaltered by TSG101 knockdown.

**Figure 4.**
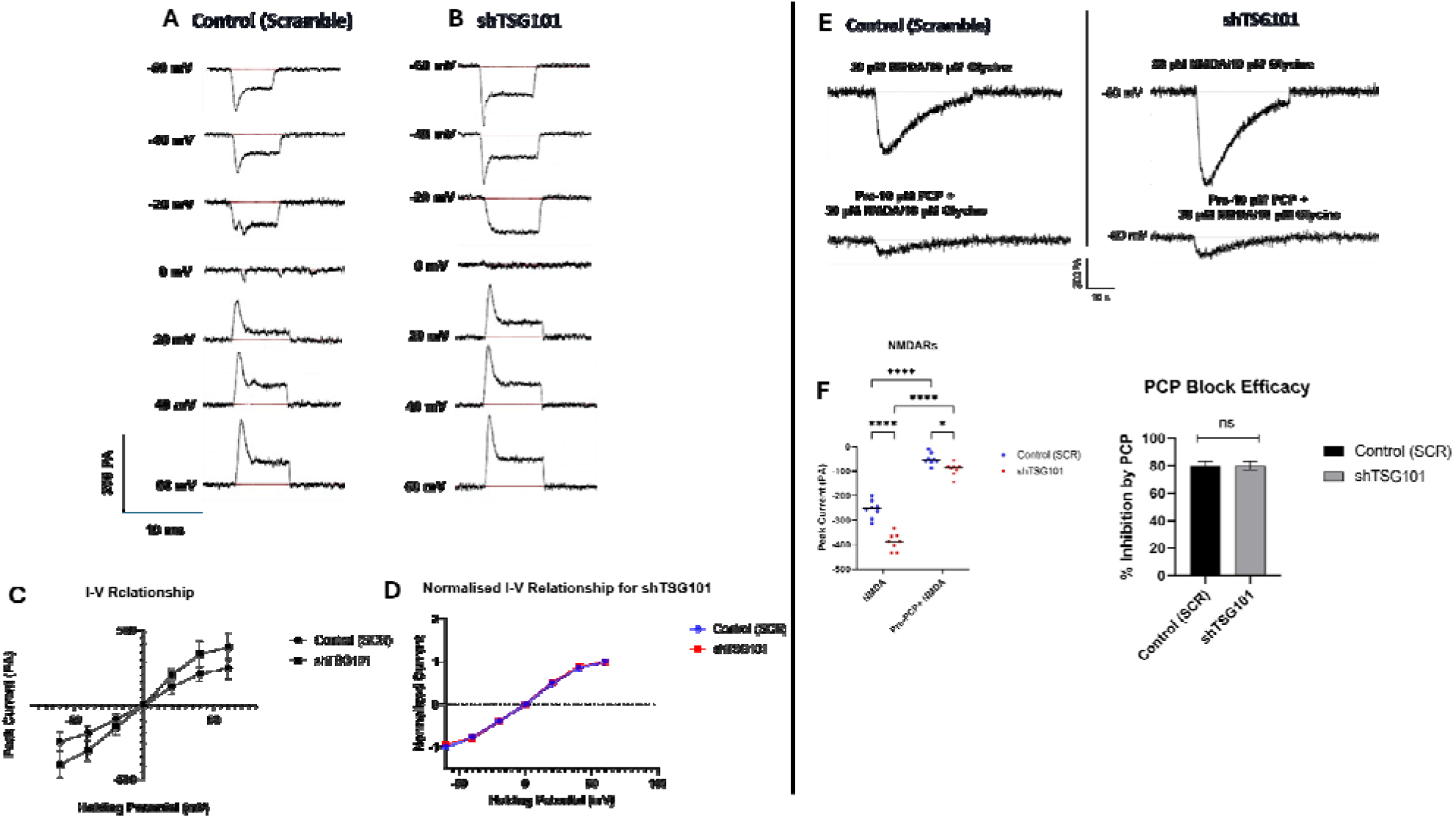
TSG101 knockdown increases NMDAR current amplitude without altering voltage-dependence or channel block. (A–B) Representative NMDAR current traces from control and shTSG101 neurons recorded at the indicated holding potentials. (C) Current–voltage (I-V) relationship demonstrating increased current amplitude in shTSG101 neurons, with no shift in the reversal potential (n = 9–11 cells per group). (D) Normalised I-V curves (normalised at +60 mV), showing identical voltage-dependence between groups. (E) Representative traces of NMDA receptor-mediated currents in the presence of PCP (10 μM) at –60 mV. (F) Peak NMDA-evoked currents (pA) in control (SCR) and shTSG101 neurons before and after PCP (10 µM) block (left) and quantification of percentage inhibition by PCP (right). Baseline currents were larger in shTSG101 than control (****p < 0.0001); PCP significantly reduced currents in both groups (****p < 0.0001 each), though a smaller group difference persisted post-block (*p < 0.05) (two-way ANOVA with Tukey’s multiple comparisons test). Percentage inhibition by PCP was equivalent between control and shTSG101 (80.2 ± 3.1% vs. 79.5 ± 2.8%; p > 0.05, ns; unpaired t-test). (n = 8–10 cells per group; ns, not significant). Data are mean ± SEM. *p < 0.05, ****p < 0.0001, ns = not significant.

To distinguish whether this increased current reflected altered channel kinetics or an expanded surface receptor pool, we examined sensitivity to the open-channel blocker PCP. Pre-application of 10 µM PCP significantly inhibited NMDAR currents in both control and shTSG101 neurons compared to baseline (*p < 0.05 for both; Fig. 4F). However, the extent of inhibition by PCP was equivalent between groups (80.2 ± 3.1% vs. 79.5 ± 2.8%; p > 0.05; ns), indicating that channel pore properties were not affected. Representative traces are shown in Fig. 4E.Together, these data demonstrate that TSG101 knockdown increases the abundance of functional surface NMDARs without altering their intrinsic biophysical or pharmacological properties.

ESCRT-I complex regulates the endosomal sorting of multiple ubiquitinated membrane proteins. We therefore, examined if TSG101 knockdown similarly affects GABA_A_ receptor membrane expression and function. Analysis of GABA_A_ receptor function revealed a parallel enhancement. shTSG101 significantly increased the peak amplitude of GABA-evoked currents across a range of concentrations (Fig. 5A-B). Specifically, the maximal current (I_max_) increased from -413 pA in controls to -770 pA in shTSG101 neurons (p < 0.01; Fig. 5C), while the EC_50_ for GABA remained unchanged (Control: 0.056 µM; shTSG101: 0.059 µM; Fig. 5D), indicating that TSG101 knockdown increases receptor number without altering agonist potency. The Hill slopes were also comparable (1.1 vs. 1.2), indicating unaltered receptor cooperativity.

**Figure 5.**
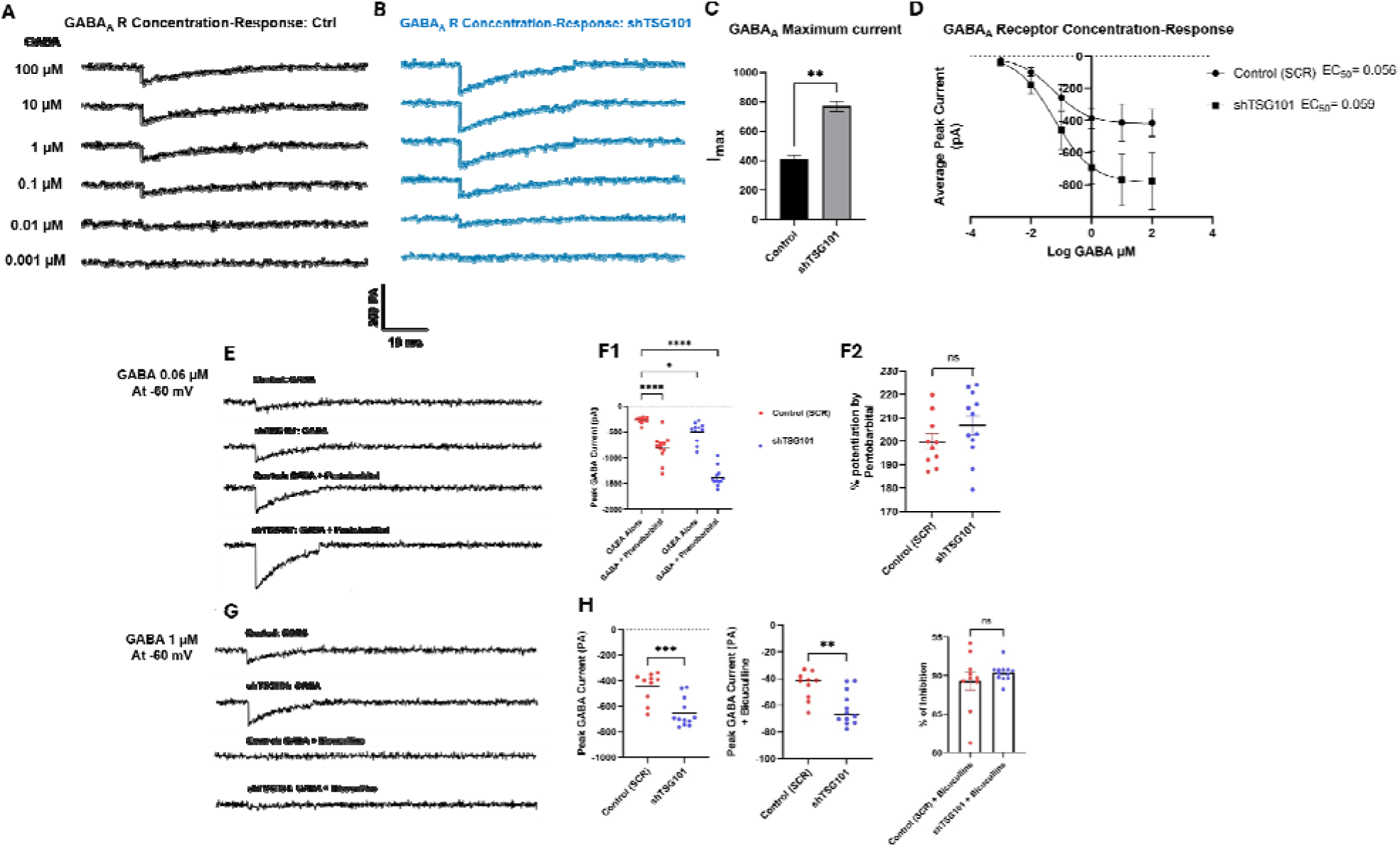
TSG101 knockdown increases GABA_A_ receptor current amplitude without altering pharmacological properties. (A–D) Concentration-response analysis of GABA_A_R-mediated currents in control (SCR) and shTSG101 neurons. (A–B) Representative traces of inward currents evoked by increasing GABA concentrations (0.001-100 µM) in control (A) and shTSG101 (B) neurons. (C) The maximal current (I_max_) for GABA_A_ increased from -413 pA in controls to -770 pA in shTSG101 neurons (p < 0.01; unpaired t-test; n = 8– 10 cells per group). (D) GABA concentration–response curves. TSG101 knockdown significantly increases maximal GABA-evoked current amplitude without altering GABA potency, as indicated by comparable EC_50_ values (Control EC_50_ = 0.056 μM; shTSG101 EC_50_= 0.059 μM; n = 8–10 cells per group). The low EC_50_ is consistent with the presence of high affinity extrasynaptic GABA_A_ receptors in cultured hippocampal neurons. (E-F) Allosteric modulation of GABA_A_Rs by pentobarbital. (E) Representative traces of GABA-evoked currents (0.06 µM, approximately EC_50_) in the absence and presence of pentobarbital (100 µM). (F) Quantification of peak GABA currents (left), showing potentiation by pentobarbital in both groups, and percentage potentiation (right), which was comparable between groups (Control: 200 ± 15%; shTSG101: 190 ± 12%; p > 0.05; n = 8–12 cells per group), indicating preserved positive allosteric modulation. (G-H) GABA_A_ receptor blockade by bicuculline. (G) Representative traces showing inhibition of GABA-evoked currents (1 µM GABA) by bicuculline (10 µM). (H) Quantification of peak currents. TSG101 knockdown significantly reduced baseline and bicuculline-sensitive current amplitudes (left and middle panels), whereas the percentage inhibition by bicuculline (right panel) was unchanged, (approximately 90% in both groups; n = 10–12 cells per group), indicating preserved antagonist sensitivity. Data are mean ± SEM. Individual data points represent three independent cultures. Statistical significance was determined using unpaired Student’s t-tests (C, F left, H left/middle) or one-way ANOVA (F right, H right). *p < 0.05, **p < 0.01, ***p < 0.001, ****p < 0.0001; ns, not significant.

Importantly, the functional integrity of GABA_A_ receptors was preserved following TSG101 knockdown. The positive allosteric modulator pentobarbital (100 µM), a barbiturate that enhances GABA_A_ receptor function by prolonging channel open time, produced equivalent potentiation of sub-maximal GABA responses (EC_50_, 0.06 µM) in both groups (Control: 200 ± 15%; shTSG101: 190 ± 12%; p > 0.05; Fig. 5E-F). Similarly, the competitive antagonist bicuculline (10 µM) produced an identical blockade of approximately 90% in both conditions (Fig. 5G-H). These findings confirm that TSG101 knockdown increases the number of functional, pharmacologically normal GABA_A_ receptors at the neuronal surface without altering their intrinsic gating or modulatory properties.

### 3.4 – TSG101 Knockdown reduces PCP-Induced Synaptic Deficits

shTSG101 increases the functional surface expression of both NMDARs and GABA_A_Rs in naïve neurons, so we tested if this enhanced receptor availability counteracts the persistent synaptic deficits induced by sub-chronic PCP. Neurons were transduced with a scrambled control or shTSG101 lentivirus prior to PCP treatment and assessed following a 48-h washout period.

shTSG101 completely prevented the persistent NMDAR hypofunction induced by PCP. Whilst control neurons (shScramble) exhibited a sustained reduction in evoked NMDAR currents following PCP washout (74% of vehicle baseline; 185.4 ± 16.3 pA), shTSG101 neurons showed a relative functional recovery, with currents reaching 108% of vehicle baseline levels (438.5 ± 41.4 pA; p < 0.0001 vs. PCP-control; Fig. 6A–C). These data indicate that the expanded surface NMDAR pool, maintained via reduced ESCRT-I-dependent degradation, provides a functional receptor reserve capable of rapidly repopulating the synapse following the removal of pharmacological blockade.

**Figure 6.**
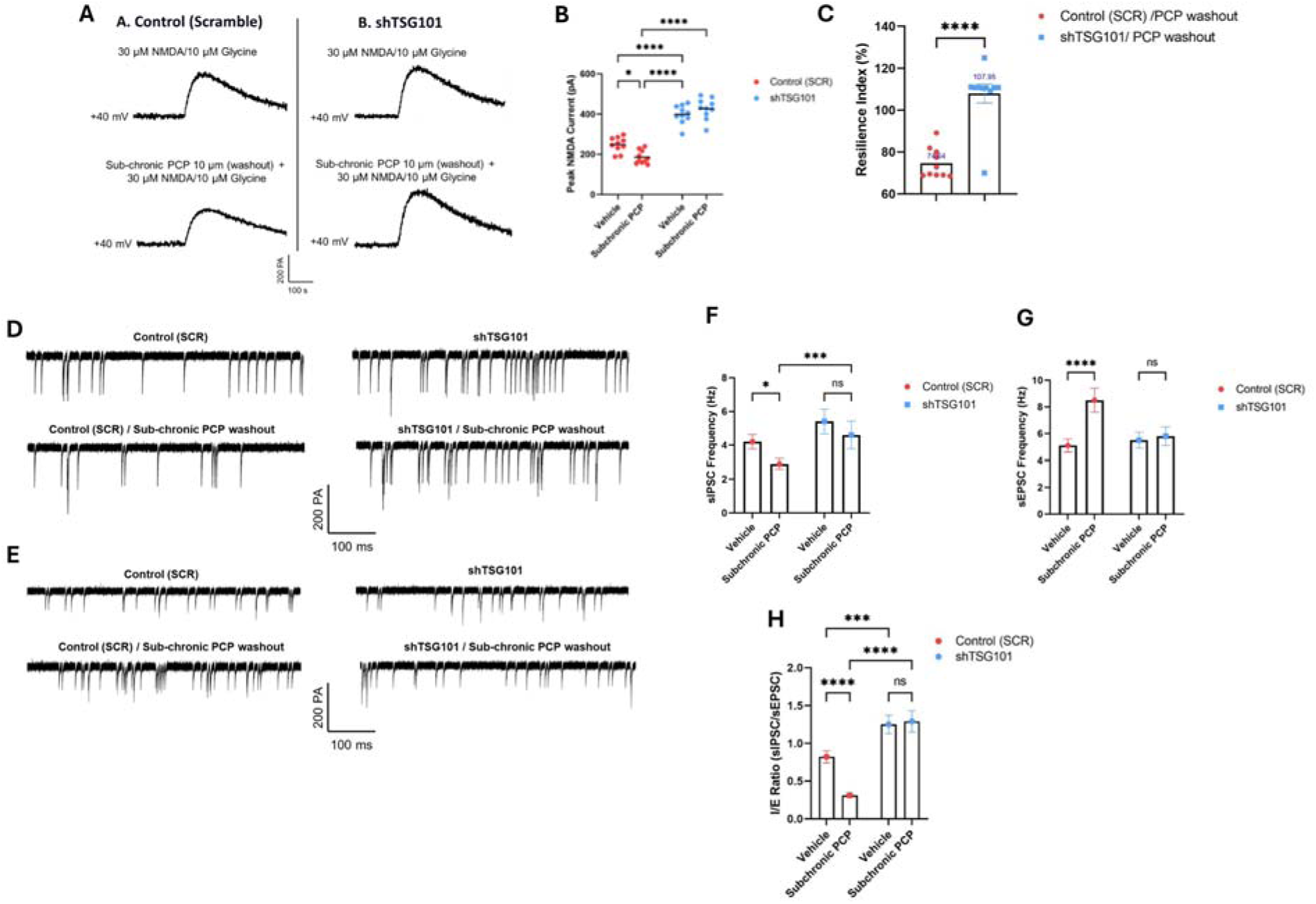
TSG101 knockdown confers resilience against PCP-induced synaptic and network deficits. (A–C) NMDA receptor-mediated currents following subchronic PCP treatment. (A) Representative whole-cell voltage-clamp recordings of evoked NMDAR-mediated currents from control (SCR) and shTSG101 neurons under vehicle and sub-chronic PCP conditions (washout). Recordings were obtained at +40 mV in Mg²⁺-free external solution containing CNQX to block AMPA receptors. Scale bar: 200 pA, 50 ms. (B) Quantification of peak NMDA current amplitudes. PCP significantly reduces NMDA currents in control neurons, whereas shTSG101 neurons exhibit larger baseline responses and are resistant to PCP-induced suppression. (C) Resilience index, calculated as the percentage of NMDA current recovered post-washout relative to the PCP-induced deficit, demonstrating significantly enhanced resilience in shTSG101 neurons compared with controls. (D&F) Spontaneous inhibitory postsynaptic currents (sIPSCs). (D) Representative traces of sIPSCs recorded at −60 mV from control and shTSG101 neurons under vehicle and PCP conditions. Scale bars: 200 pA, 100 ms. (F) Quantification of sIPSC frequency. PCP significantly reduces inhibitory event frequency in control neurons, whereas shTSG101 neurons maintain elevated inhibitory activity that is unaffected by PCP treatment. (E&G) Spontaneous excitatory postsynaptic currents (sEPSCs). (E) Representative traces of sEPSCs recorded at −60 mV from control and shTSG101 neurons under vehicle and PCP conditions. Scale bars: 200 pA, 100 ms. (G) Quantification of sEPSC frequency. Subchronic PCP increases excitatory event frequency in control neurons (consistent with network disinhibition following NMDAR hypofunction), whereas shTSG101 neurons display elevated basal excitatory activity that remains stable following PCP exposure. (H) Inhibitory/Excitatory (I/E) balance, calculated as the ratio of sIPSC to sEPSC frequency. PCP treatment induces a shift in network balance in control neurons, whereas shTSG101 neurons maintain a stable I/E ratio under both vehicle and PCP conditions. These findings indicate that TSG101 depletion promotes resistance to PCP-induced synaptic dysfunction and stabilises network activity. Data are mean ± SEM; n = 10–12 cells per group from three independent neurons. For multiple group comparisons (B, E, G, H), one-way ANOVA with Tukey’s post-hoc test was used. For two-group comparisons (C), unpaired t-test was used. *P < 0.05, **P < 0.01, ***P < 0.001, ****P < 0.0001; ns, not significant.

Analysis of inhibitory transmission revealed a parallel pattern of resilience. Control neurons showed persistent impairment of GABAergic transmission after PCP washout, with sIPSC frequency remaining significantly below the vehicle baseline (69% of baseline; 2.9 ± 0.3 Hz). In contrast, shTSG101 neurons exhibited complete recovery of sIPSC frequency to 109% of baseline (4.6 ± 0.4 Hz; p < 0.001 vs. PCP-control; Fig. 6D–F). Consistent with this restored inhibitory tone, the PCP-induced elevation in sEPSC frequency was fully normalised in shTSG101 neurons (105% of baseline; 5.4 ± 0.5 Hz), whilst control networks remained hyperexcitable (167% of baseline; 8.5 ± 0.8 Hz; p < 0.01 vs. PCP-shTSG101; Fig. 6G).

The net effect on network homeostasis was quantified by the I/E balance ratio (sIPSC/sEPSC frequency). In control neurons, PCP washout reduced the I/E ratio from 0.82 ± 0.08 to 0.31 ± 0.04 (p < 0.001; Fig. 6H), indicating a persistent shift to-wards network disinhibition. In contrast, shTSG101 neurons maintained a stable I/E ratio across vehicle and PCP conditions (1.25 ± 0.12 vs. 1.29 ± 0.13; p > 0.05), demonstrating resilience against NMDAR blockade. The higher baseline I/E ratio in shTSG101 neurons reflects the increased surface expression of both excitatory and inhibitory receptors observed in naïve neurons (Section 3.3), consistent with a home-ostatic rebalancing of network tone. Because sEPSCs and sIPSCs were recorded from the same population of pyramidal-like neurons, these data show that shTSG101 stabilises the net balance of excitatory and inhibitory inputs converging onto pyramidal neurons. However, our data do not distinguish whether this stabilisation reflects cell-type-specific receptor upregulation (e.g., NMDARs on interneurons; GABA_A_Rs on pyramidal neurons) or a general scaling of both receptor types across all neurons. Importantly, because shTSG101 was expressed prior to the PCP challenge in these experiments, these findings demonstrate a resilience phenotype.

### 3.5 – ESCRT Modulation Bidirectionally Regulates NMDAR Sensitivity to PCP in Primary Neurons

To complement and extend the electrophysiological characterisation of PCP-induced NMDAR hypofunction, we examined whether sub-chronic ESCRT modulation alters NMDAR functional sensitivity at the population level using calcium imaging. Cultures were treated with the same sub-chronic PCP protocol (0.1–30 µM, 7 days, followed by 48 h washout) used in Section 3.2, after which NMDA-evoked intracellular calcium responses were quantified. In control neurons, NMDA application elicited a rapid increase in intracellular calcium followed by a sustained plateau, consistent with prolonged receptor activation (Fig. 7C). PCP produced a concentration-dependent suppression of this response, significantly reducing both peak amplitude (ΔF/F₀) and the area under the curve (AUC), with near-complete inhibition achieved at 30 µM (Fig. 7D-F). These effects reflect a suppression of both the transient and sustained components of NMDAR-mediated calcium signalling.

**Figure 7.**
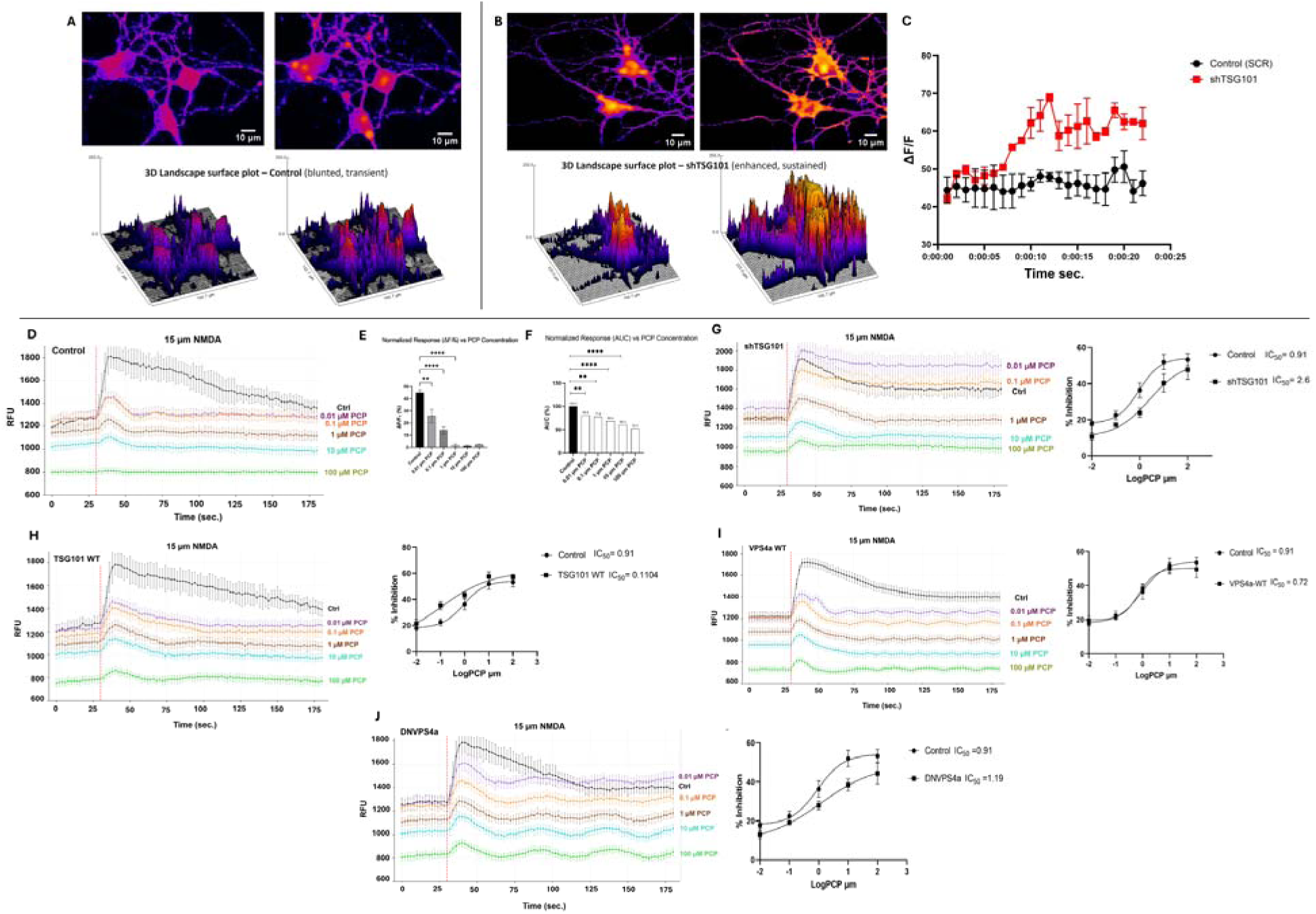
ESCRT modulation bidirectionally regulates NMDAR-mediated calcium dynamics and pharmacological sensitivity in primary neurons. (A–C) Live-cell calcium imaging of NMDA-evoked responses. (A-B) Representative fluorescence images and 3D intensity surface plots from control (A) and shTSG101 (B) neurons before and after NMDA stimulation, showing enhanced and sustained calcium responses in shTSG101 neurons. Scale bars: 10 μm. (C) Quantification of fluorescence changes (ΔF/F) over time, demostrating that TSG101 knockdown significantly increases the magnitude and duration of calcium responses. (D–F) Concentration-dependent inhibition of NMDA-evoked calcium responses by PCP in control neurons. (D) Representative time-course traces showing NMDA responses (15 µM) in the presence of increasing PCP concentrations (0.01–100 µM). (E-F) Quantification of normalized response amplitudes (ΔF/F₀) (E) and area under the curve (AUC) (F), showing progressive, dose-dependent suppression of NMDA responses by PCP (**p < 0.01, ****p < 0.0001; one-way ANOVA with Dunnett’s post-hoc test; n = 4–6 independent experiments). (G-J) Pharmacological sensitivity of NMDA-evoked calcium responses following ESCRT modulation. (G) Concentration–inhibition curve in shTSG101 neurons, revealing a rightward shift in PCP sensitivity compared to controls (Control IC_50_ = 0.91 μM; shTSG101 IC_50_ = 2.6 μM). (H) Overexpression of wild-type (WT) TSG101 increases PCP potency, producing a leftward shift in the concentration–inhibition relationship (Control IC_50_ = 0.91 μM; TSG101 WT IC_50_ = 0.1104 μM). (I) Neurons expressing WT-VPS4a exhibit PCP sensitivity comparable to control (IC_50_ = 0.72 μM). (J) Expression of dominant-negative VPS4a (DNVPS4a) partially attenuates PCP inhibition, shifting the concentration–response relationship rightward (IC_50_ = 1.19 μM). Data are mean ± SEM from independent neuronal cultures or cells. Statistical significance was determined using one-way or two-way ANOVA with appropriate post-hoc multiple-comparison testing. *P < 0.05, **P < 0.01, ***P < 0.001, ****P < 0.0001.

To visualise the spatial distribution and magnitude of these calcium signals, we generated 3D surface landscapes of calcium intensity (ΔF/F₀) over space. Control neurons exhibited blunted, transient calcium elevations with rapid signal decay (Fig. 7A), whilst shTSG101 neurons displayed markedly enhanced and sustained elevations with greater peak amplitudes (Fig. 7B). These data provide a spatial representation consistent with an expanded functional surface NMDAR pool. Temporal dynamics confirmed that TSG101 knockdown resulted in a prolonged elevation of intracellular calcium following NMDA stimulation relative to scrambled controls (Fig. 7C).

TSG101 knockdown (shTSG101) enhanced calcium signals and produced a rightward shift in the PCP concentration–response curve; the IC_50_ increased from 0.91 µM in controls to 2.6 µM in shTSG101 neurons, representing an approximately 2.9-fold reduction in PCP potency (p < 0.05; Fig. 7G). Again, this finding is consistent with an expanded surface NMDAR pool that requires higher antagonist concentrations to achieve equivalent blockade. These observations were further corroborated by live-cell calcium imaging (Supplementary Videos 1–2), in which shTSG101-expressing neurons exhibited markedly enhanced and sustained responses to NMDA stimulation compared with controls, providing direct visual confirmation of the quantitative findings.

TSG101 overexpression reduced NMDA-evoked calcium responses and increased PCP potency, with the IC_50_ decreasing from 0.91 µM to 0.39 µM (a 2.3-fold increase in potency; p < 0.05; Fig. 7H). These data are consistent with a depleted surface NMDAR population resulting from enhanced ESCRT-I-mediated receptor degradation.

Perturbation of ESCRT-III function via DN-VPS4a produced an effect that was qualitatively similar to, but quantitatively more modest than, shTSG101, inducing a rightward shift in PCP sensitivity (IC_50_: 1.19 µM; p < 0.05; Fig. 7I), indicative of increased functional receptors. VPS4a overexpression produced the opposite pattern, shifting PCP sensitivity leftward (IC_50_: 0.72 µM; p < 0.05; Fig. 7J), consistent with enhanced ESCRT-III-mediated membrane scission and receptor turnover. Across all conditions, changes in pharmacological sensitivity closely tracked alterations in response amplitude and duration, confirming that ESCRT activity bidirectionally regulates the functional surface availability of NMDARs by balancing endosomal receptor degradation against recycling back to the plasma membrane.

### 3.6 – TSG101 Knockdown Rescues Molecular Hallmarks of Glutamatergic Dysfunction

We next examined the effects of shTSG101 on synaptic and signalling proteins disrupted by sub-chronic PCP. Sub-chronic PCP treatment significantly reduced both PSD-95 puncta density and BDNF fluorescence intensity in primary neurons (Fig. 8A–B, D; p < 0.05 and p < 0.001, respectively). Expression of shTSG101 significantly reversed both deficits, restoring PSD-95 and BDNF levels towards vehicle control values (Fig. 8C, E). The restoration by shTSG101 of sub-chronic PCP-induced PSD-95 is consistent with the electrophysiological recovery of NMDAR currents detailed in Section 3.4. In contrast, DN-VPS4a failed to rescue either PSD-95 or BDNF, and wild-type TSG101 overexpression (TSG101-OE) did not counteract PCP-induced reductions (Fig. 8A, E). The inability of DN-VPS4a to rescue these molecular markers, despite its partial effect on surface receptor levels, suggests that complete functional recovery requires the specific relief of ESCRT-I-mediated cargo sorting rather than broader perturbation of ESCRT-III-dependent membrane scission.

**Figure 8.**
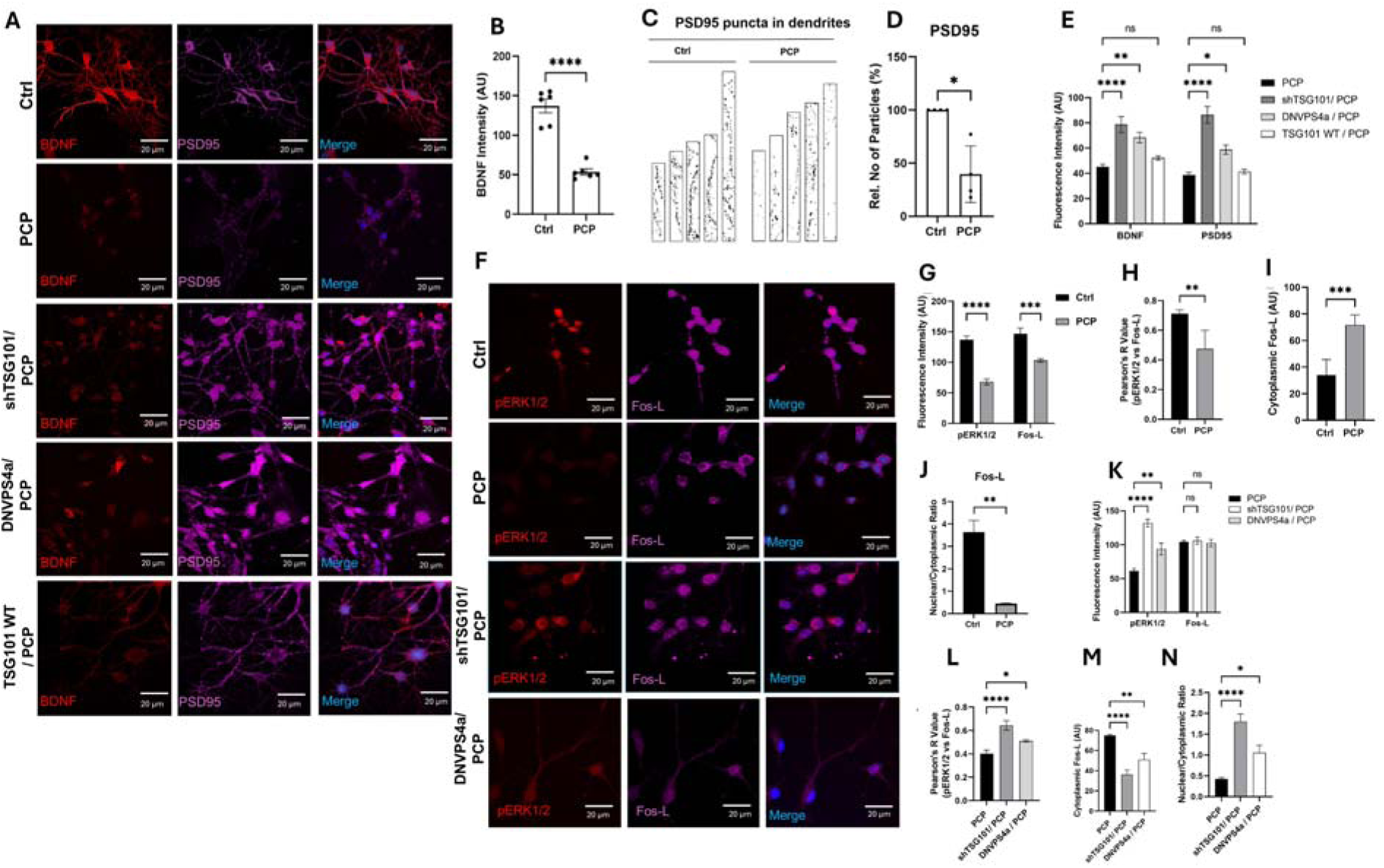
TSG101 knockdown rescues PCP-induced deficits in synaptic protein expression, ERK1/2 signalling, and Fos-L nuclear translocation. (A-D) Effect of subchronic PCP treatment on BDNF and PSD-95 expression. (A) Representative immunofluorescence images of BDNF (red) and PSD-95 (magenta). (B) Quantification of BDNF fluorescence intensity (***p < 0.001). (C) Representative images of dendritic PSD-95 puncta. (D) Quantification of PSD-95 puncta density (*p < 0.05), expressed as percentage of control. Scale bar: 20 µm. (E) Quantification of BDNF and PSD-95 intensity across experimental groups (PCP, shTSG101/PCP, DN-VPS4a/PCP, and TSG101-OE/PCP). shTSG101 rescues PCP-induced deficits, whereas DN-VPS4a or TSG101-OE do not (one-way ANOVA with Tukey’s post-hoc test). (F-J) Effect of PCP treatment on pERK1/2 and Fos-L signaling. (F) Representative images of pERK1/2 (red) and Fos-L (magenta). (G) Quantification of pERK1/2 and Fos-L fluorescence intensity. (H) Pearson’s correlation coefficient (*R*) for pERK1/2–Fos-L colocalization. (I-J) Cytoplasmic Fos-L intensity (I) and nuclear/cytoplasmic Fos-L ratio (J). (K-N) Rescue of pERK1/2 and Fos-L signaling by ESCRT modulation. Quantification across PCP, shTSG101/PCP, and DN-VPS4a/PCP conditions for pERK1/2 and Fos-L intensity (K), pERK1/2–Fos-L colocalisation (L), cytoplasmic Fos-L intensity (M), and nuclear/cytoplasmic Fos-L ratio (N). shTSG101 promotes recovery of signaling markers, whereas DN-VPS4a does not (one-way ANOVA with Tukey’s post-hoc test). Data are mean ± SEM. Significance levels: *p < 0.05, **p < 0.01, ***p < 0.001, ****p < 0.0001; ns, not significant.

Phosphorylation of ERK1/2 at Thr202/Tyr204 (pERK1/2), a marker of activity-dependent intracellular signalling, was significantly reduced by sub-chronic PCP (Fig. 8F–G; p < 0.0001). shTSG101 fully restored pERK1/2 fluorescence intensity to vehicle control levels (Fig. 8K), whilst DN-VPS4a again failed to rescue the deficit, further distinguishing the molecular consequences of ESCRT-I versus ESCRT-III perturbation.

The subcellular localisation of the immediate early gene product Fos-L was examined next as a readout of activity-dependent transcriptional activation. Sub-chronic PCP induced pronounced cytoplasmic retention of Fos-L, reflected by a significant increase in cytoplasmic intensity (Fig. 8I, M; p < 0.001) and a reduced nuclear-to-cytoplasmic ratio (Fig. 8J, N; p < 0.01), indicating impaired activity-dependent nuclear import. shTSG101 restored the nuclear translocation of Fos-L, significantly increasing the nuclear/cytoplasmic ratio and reducing cytoplasmic accumulation toward control levels (Fig. 8J, M–N; p < 0.0001), whilst DN-VPS4a produced no significant rescue.

Finally, the functional coupling between ERK1/2 activation and Fos-L induction was assessed by quantifying pERK1/2–Fos-L colocalisation. Sub-chronic PCP significantly reduced Pearson’s correlation coefficients for pERK1/2 and Fos-L (Fig. 8H; p < 0.01), indicative of disrupted ERK-to-transcription factor signalling. shTSG101 restored this colocalisation to control levels (Fig. 8L), whilst DN-VPS4a showed no significant effect (Fig. 8L), consistent with its failure to rescue upstream pERK1/2 activation.

Together, these findings demonstrate that TSG101 knockdown counteracts PCP-induced deficits across all major molecular readouts examined – including synaptic scaffolding (PSD-95), neurotrophic support (BDNF), ERK1/2 activation, Fos-L nuclear translocation, and ERK–Fos-L functional coupling – thereby reactivating the transcriptional and synaptic plasticity programmes required for long-term, activity-dependent neuronal adaptation.

## 4 – Discussion

In this study, we demonstrate that inhibition of the ESCRT-I component TSG101 confers protects against synaptic and network deficits induced by sub-chronic NMDAR blockade with PCP, a validated pharmacological model of schizophrenia-relevant pathology [1, 2]. shTSG101 enhances the surface expression of functional NMDARs, and GABA_A_Rs without altering their intrinsic biophysical properties. Rather than selectively enhancing excitation or inhibition, TSG101 knockdown increases the surface availability of both receptors’ classes.

By upregulating both receptor classes, this counters the core deficit induced by PCP, NMDAR hypofunction, which preferentially impairs NMDAR function on GABAergic interneurons, reducing their firing and GABA release, and leading to network disinhibition. By upregulating NMDARs on interneurons, shTSG101 restores their excitability and GABA release, while concurrent upregulation of GABA_A_Rs on pyramidal neurons enhances their sensitivity to the restored inhibition. This coordinated action enables recovery of neurotransmission and stabilises the excitation/inhibition (E/I) balance following pathological challenge. Consistent with this, shTSG101 prevented the PCP-induced loss of inhibitory transmission, normalised network excitatory activity, and reactivates ERK-dependent transcriptional pathways associated with synaptic plasticity. Together, these findings identify ESCRT-I-dependent receptor trafficking as a regulator of synaptic homeostasis and suggest that limiting receptor degradation may represent a mechanism-based strategy for disorders characterised by glutamatergic hypofunction. The conclusion that TSG101 knockdown increases surface receptor abundance, rather than altering channel properties, rests on convergent pharmacological evidence. shTSG101 increased the amplitude of evoked NMDAR and GABA_A_R currents without shifting voltage-dependence, reversal potentials, agonist potencies, Hill slopes, or sensitivities to allosteric modulators and competitive antagonists, a pharmacological signature indicative of an expanded surface receptor pool [19, 20]. Furthermore, the preserved I-V relationship and equivalent PCP-mediated blockade in both shTSG101 and control neurons confirm that the fundamental biophysical properties of NMDARs remain entirely intact.

Mechanistically, these effects align with the established role of ESCRT-I complex in sorting early endosomal cargo [29]. TSG101 recognises ubiquitinated membrane proteins and facilitates their incorporation into the intraluminal vesicles of multivesicular bodies targeted for lysosomal degradation [24, 27]. Consequently, a loss of TSG101 function is expected to impair this degradative sorting pathway, causing internalised receptors to accumulate within early endosomal compartments and increasing their probability of recycling back to the plasma membrane [48]. Although this model requires direct confirmation via high-resolution, live-cell endosomal imaging in future work, the proposed “receptor reservoir” mechanism provides a compelling explanation for this functional recovery. Specifically, this mechanism predicts that internalised receptors accumulate in early endosomal compartments following TSG101 knockdown, enabling rapid recycling to the surface upon PCP wash-out, whereas control neurons would rely on slower de novo synthesis. Direct testing of this prediction—for example, by quantifying endosomal accumulation of NMDARs using organelle-specific markers—would be valuable to confirm this mechanism.

The concurrent upregulation of both NMDAR and GABA_A_R surface expression following shTSG101 knockdown is mechanistically pivotal for understanding the restoration of network I/E balance. Cortical disinhibition in schizophrenia is thought to arise primarily from NMDAR hypofunction on parvalbumin-positive GABAergic interneurons, which subsequently diminishes their inhibitory output onto pyramidal neurons [9, 10]. Although NMDAR ‘hypofunction’ might intuitively suggest a global suppression of network activity, the preferential vulnerability of interneurons to NMDAR blockade produces a net increase in pyramidal neuron excitability and a collapse of E/I balance. Our observation that sub-chronic PCP reduces sIPSC frequency while increasing sEPSC frequency (Fig. 3) is entirely consistent with this disinhibition model.

By enhancing the surface availability of both NMDARs and GABA_A_Rs, shTSG101 could, in principle, restore network balance through two complementary mechanisms: (i) increased NMDAR availability on interneurons may restore their excitability and GABA release, and (ii) increased GABA_A_R availability on pyramidal neurons may enhance their sensitivity to the restored inhibition. However, our current data, obtained from mixed cortical cultures, do not distinguish between these possibilities, as we cannot determine the cell-type-specific distribution of receptor upregulation. Both sEPSCs and sIPSCs were recorded from pyramidal-like neurons, allowing assessment of the net balance of excitation and inhibition converging onto the same cell type. The observation that shTSG101 upregulates both NMDARs and GABA_A_Rs in mixed cultures is consistent with both cell-type-specific and general scaling mechanisms. Nevertheless, the net effect at the network level is clear: shTSG101 prevented the PCP-induced loss of inhibitory tone and stabilised the I/E ratio (Fig. 6). Future studies using cell-type-specific manipulations will be required to dissect the relative contributions of interneuron NMDAR upregulation versus pyramidal neuron GABA_A_R upregulation to the observed resilience phenotype. This coordinated action across both neurotransmitter receptor systems distinguishes ESCRT-I inhibition from conventional strategies that exclusively target either the excitatory arm (e.g., glycine-site agonists or D-serine) or the inhibitory arm (e.g., benzodiazepines or GABA reuptake inhibitors) of the E/I balance [14, 15, 49].

Our finding that GABABR1 surface levels were selectively increased by TSG101 knockdown, but not by DN-VPS4a, suggests that trafficking of this GPCR is preferen-tially regulated by ESCRT-I rather than ESCRT-III. Given that GABAB receptors mod-ulate both presynaptic neurotransmitter release and postsynaptic membrane excita-bility, their disruption represents a core component of the broader GABAergic deficits observed in schizophrenia pathophysiology [50]. Whether this selectivity extends to other GPCRs, including mGluRs, remains to be determined. The concept of network resilience is central to interpreting the functional outcomes of TSG101 knockdown. shTSG101 neurons maintained a stable I/E balance under both baseline conditions and following the PCP challenge, whilst control networks exhibited a sustained collapse of the I/E ratio. This resilience is driven by the enhanced inhibitory tone conferred by increased GABA_A_R and GABA_B_R surface expression, which provides a homeostatic buffer against excitatory overshoot. Specifically, the recovery of sIPSC frequency to 109% of baseline following PCP washout in shTSG101 neurons suggests that enhanced GABAergic transmission actively constrains the hyperexcitability that would otherwise emerge. This finding is highly consistent with literature demonstrating that strengthening inhibitory circuits prevents pathological network disinhibition [11, 49].

The elevated baseline I/E ratio in shTSG101 neurons did not result in hypo-excitability, as sEPSC frequency remained stable (Fig. 6G). This suggests homeostatic regulation constrains excessive inhibition. The elevated I/E ratio reflects the increased surface availability of both excitatory and inhibitory receptors (Figs. 1–2), indicating a rebalancing of network tone rather than non-specific hyperinhibition. This functional rescue extended beyond the cell surface to synaptic architecture and downstream intracellular cascades. shTSG101 effectively restored both PSD-95 puncta density and BDNF fluorescence intensity, which were substantially depleted following sub-chronic PCP exposure. The recovery of PSD-95 is mechanistically aligned with our electrophysiological rescue profile, as this core scaffolding protein anchors NMDARs at postsynaptic sites and structurally links them to downstream signalling complexes [16, 17]. Concurrently, BDNF signalling through the TrkB receptor, is known to activate the ERK1/2 cascade, thereby supporting NMDAR surface expression and synaptic stabilisation [51, 52]. These interconnected interactions provide a plausible mechanistic sequence wherein shTSG101-mediated restoration of BDNF drives ERK1/2 reactivation, which subsequently promotes nuclear Fos-L translocation and activity-dependent gene expression.

Notably, Fos-L (ΔFosB) is a uniquely stable member of the AP-1 transcription factor family with an exceptionally long half-life spanning days to weeks [53], allowing it to regulate genes involved in synaptic plasticity and long-term neuronal adaptation. Its complete restoration by shTSG101 suggests that ESCRT-I inhibition may induce persistent transcriptional changes capable of supporting sustained synaptic recovery. Crucially, the failure of DN-VPS4a to rescue any of these molecular markers — despite its partial effect on surface receptor levels — indicates that ESCRT-III perturbation alone is insufficient to engage the full molecular programme of recovery. Instead, complete reversal rescue requires the specific relief of ESCRT-I-mediated cargo commitment to the degradative pathway. Based on these findings, we propose an integrated model whereby the inhibition of ESCRT-I-dependent receptor degradation expands surface receptor availability, enhances PSD95-BDNF-ERK1/2 signalling, and ultimately preserves network stability during states of chronic NMDAR hypofunction (Fig. 9).

**Figure 9.**
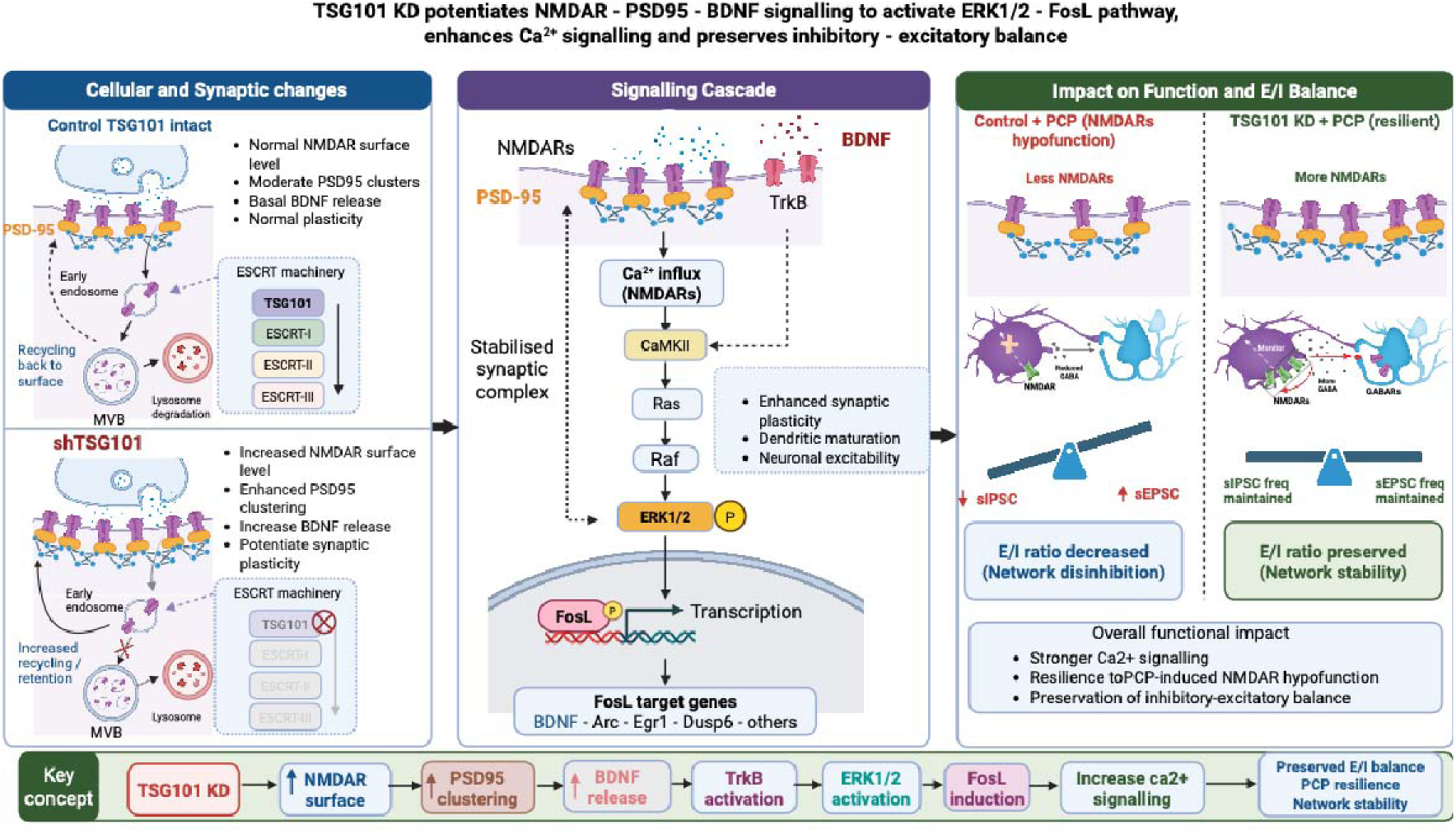
TSG101 knockdown enhances NMDA receptor signalling and preserves excitatory–inhibitory balance under PCP-induced hypofunction. **Left panel: Cellular and synaptic changes.** Under physiological conditions (Control), intact TSG101-mediated ESCRT function regulates NMDA receptor (NMDAR) trafficking through endosomal sorting, multivesicular body (MVB) formation, recycling, and lysosomal degradation, maintaining homeostatic receptor surface expression, PSD-95 clustering, basal BDNF release, and synaptic plasticity. Conversely, TSG101 knockdown (shTSG101) disrupts ESCRT-dependent sorting, reducing lysosomal degradation and promoting receptor retention and recycling to the plasma membrane. This leads to elevated surface NMDAR expression, enhanced PSD-95 clustering, increased BDNF release, and potentiated synaptic signalling. **Middle panel: Signalling cascade.** Increased surface NMDAR availability enhances intracellular Ca²⁺ influx upon receptor activation. Elevated Ca²⁺ stimulates the CaMKII–Ras–Raf cascade, driving phosphorylation and activation of ERK1/2. Concurrently, increased BDNF release activates TrkB receptors, further potentiating ERK1/2 phosphorylation. Activated ERK1/2 induces the transcription factor FosL, promoting the transcription of activity-dependent target genes including BDNF, Arc, Egr1, and Dusp6. These molecular adaptations support synaptic strengthening, dendritic maturation, and neuronal resilience. **Right panel: Impact on Function and E/I Balance.** In control neurons, sub-chronic PCP treatment blocks NMDARs, suppressing glutamatergic transmission, reducing inhibitory synaptic activity (sIPSC frequency), elevating excitatory drive (sEPSC frequency), and disrupting the excitatory–inhibitory (E/I) balance to cause network disinhibition. Conversely, shTSG101 neurons exhibit elevated surface NMDAR availability and robust downstream signalling, offsetting PCP-induced receptor blockade. This compensatory mechanism sustains inhibitory and excitatory synaptic activity, thereby maintaining the E/I ratio and network stability. **Overall model.** TSG101 depletion increases NMDAR surface density and PSD-95 stabilization, activates BDNF–TrkB and ERK1/2–FosL pathways, and amplifies Ca²⁺-dependent synaptic transmission. These coordinated adaptations confer resilience against PCP-induced NMDAR hypofunction, preserving E/I balance and neuronal network stability. **Abbreviations:** Arc, activity-regulated cytoskeleton-associated protein; BDNF, brain-derived neurotrophic factor; CaMKII, calcium/calmodulin-dependent protein kinase II; Dusp6, dual specificity phosphatase 6; E/I, excitatory/inhibitory; Egr1, early growth response 1; ERK1/2, extracellular signal-regulated kinase 1/2; ESCRT, endosomal sorting complex required for transport; FosL, Fos-like transcription factor; MVB, multivesicular body; NMDAR, N-methyl-D-aspartate receptor; PCP, phencyclidine; PSD-95, postsynaptic density protein 95; sEPSC, spontaneous excitatory postsynaptic current; sIPSC, spontaneous inhibitory postsynaptic current; shTSG101, short hairpin RNA-targeting TSG101; TrkB, tropomyosin receptor kinase B. *Note: This schematic illustrates one hypothesised mechanism by which shTSG101 could restore E/I balance: NMDAR upregulation on interneurons (restoring their excitability) and GABA∼A∼R upregulation on pyramidal neurons (enhancing postsynaptic inhibition). However, our current data from mixed cortical cultures do not resolve the cell-type-specific distribution of receptor upregulation. The net stabilisation of the I/E ratio is clearly demonstrated, but the relative contributions of each cell type remain to be tested using cell-type-specific manipulations in future studies*.

The therapeutic potential of ESCRT modulation lies in its ability to address the underlying trafficking deficits that drive to NMDAR hypofunction. Current strategies for targeting glutamatergic hypofunction focus on directly potentiating NMDAR activity through glycine-site agonists, D-serine, or positive allosteric modulators [14, 15], or on modulating upstream signalling via mGluR2/3 agonists or muscarinic M1/M4 receptor agonists [54, 55]. Whilst these strategies can enhance receptor function acutely, they do not correct the fundamental abnormalities in receptor trafficking and surface stability thought to characterise schizophrenia pathophysiology [16, 56]. By preventing lysosomal degradation of internalised receptors, ESCRT-I inhibition establishes a functional receptor reserve that can be mobilised rapidly upon demand. This mechanism is exemplified by the complete recovery of NMDAR currents following PCP washout in shTSG101 neurons. Our calcium imaging data further substantiate this interpretation, demonstrating that shTSG101 expression reduces the potency of NMDAR channel blockers — a finding consistent with an expanded surface receptor pool requiring higher antagonist concentrations for equivalent occupancy. Collectively, these data suggest that targeting endosomal receptor fate, rather than receptor activation *per se*, represent a more durable approach to restoring glutamatergic function.

The clinical plausibility of this paradigm is underscored by transcriptomic analyses of post-mortem schizophrenia tissue, which demonstrates dysregulation of multiple ESCRT complex genes, including key components of ESCRT-0 (HGS, STAM), ESCRT-I (VPS37A, TSG101), and ESCRT-III (CHMP2B, CHMP4B, CHMP6) [7, 8]. Our findings establish a mechanistic framework linking such dysregulation to NMDAR hypofunction and network instability, suggesting that therapeutic normalisation of ESCRT-I activity, via small-molecule inhibitors of TSG101-cargo interactions or gene therapy, could confer robust synaptic resilience. Importantly, this approach may offer precise target selectivity; the observation that DN-VPS4a produced overlapping yet distinct effects compared to shTSG101 across receptor systems and molecular markers raises the possibility that individual ESCRT pathway nodes could be targeted selectively depending on the receptor populations and clinical features of interest.

In conclusion, this study identifies ESCRT-I-dependent receptor trafficking as an important regulator of synaptic homeostasis and E/I balance in neurons. Genetic inhibition of TSG101 enhances the surface availability of NMDARs, and GABA_A_Rs, without altering their intrinsic biophysical properties, thereby conferring resilience against glutamatergic hypofunction at both the synaptic and network scales. These findings provide a strong rationale for investigating pharmacological ESCRT-I modulators to combat schizophrenia-relevant pathology.

## 6 – Author Contributions

M.F.S. performed all experiments, analysed data, and drafted the manuscript. V.P, J.E.G, and N.G revised electrophysiology experiments and contributed to data interpretation and revisions. S.L.M. supervised all aspects of the work and revised the manuscript. S.K. conceived the study, supervised all aspects of the work, and wrote the final manuscript. All authors reviewed and approved the final version.

## 8 – Data availability

The datasets generated and/or analysed during the current study are available from the corresponding author upon reasonable request.

## 9 – Ethics Statement

All procedures involving animals were approved by the University of Bradford Animal Welfare and Ethical Review Body (AWERB) and complied with The Animals (Scientific Procedures) Act 1986 (UK).

